# Global observation of plankton communities from space

**DOI:** 10.1101/2022.09.23.508961

**Authors:** Hiroto Kaneko, Hisashi Endo, Nicolas Henry, Cédric Berney, Frédéric Mahé, Julie Poulain, Karine Labadie, Odette Beluche, Roy El Hourany, Tara Oceans Coordinators, Samuel Chaffron, Patrick Wincker, Ryosuke Nakamura, Lee Karp-Boss, Emmanuel Boss, Chris Bowler, Colomban de Vargas, Kentaro Tomii, Hiroyuki Ogata

## Abstract

Satellite remote sensing from space is a powerful way to monitor the global dynamics of marine plankton. Previous research has focused on developing models to predict the size or taxonomic groups of phytoplankton. Here we present an approach to identify representative communities from a global plankton network that included both zooplankton and phytoplankton and using global satellite observations to predict their biogeography. Six representative plankton communities were identified from a global co-occurrence network inferred using a novel rDNA 18S V4 planetary-scale eukaryotic metabarcoding dataset. Machine learning techniques were then applied to train a model that predicted these representative communities from satellite data. The model showed an overall 67% accuracy in the prediction of the representative communities. The prediction based on 17 satellite-derived parameters showed better performance than based only on temperature and/or the concentration of chlorophyll *a*. The trained model allowed to predict the global spatiotemporal distribution of communities over 19-years. Our model exhibited strong seasonal changes in the community compositions in the subarctic-subtropical boundary regions, which were consistent with previous field observations. This network-oriented approach can easily be extended to more comprehensive models including prokaryotes as well as viruses.

## Introduction

Monitoring the global dynamics of marine plankton is essential to understand the function of marine microbial ecosystem and its interaction and evolution with climate change. It can also facilitate the discovery of new plankton species. However, it is impossible to obtain global plankton samples at high spatial and temporal density using research ships alone due to the extent of the ocean. Regular and global remote sensing using satellites can potentially be used to solve this problem. Because plankton pigments absorb light, the spectrum of light reflected from the ocean surface that is observed by satellites (remote sensing reflectance) has a specific relationship with the plankton composition. Environmental parameters such as sea surface temperature (SST) are also related with the plankton composition [1].

Several models for predicting plankton community satellite-derived data have been developed over the past decades [2, 3]. Most of them focused on phytoplankton because these species contain pigments, such as chlorophylls, carotenoids, and phycobilins, to capture light energy for photosynthesis [4]. The abundances of three size classes, micro-phytoplankton (> 20 µm), nano-phytoplankton (2–20 µm) and pico-phytoplankton (0.2–2 µm), can be predicted with simple models only integrating the concentration of chlorophyll *a* ([Chl]), which is the core of the photosynthetic unit [5–7]. More advanced models have also been developed to predict size classes using remote sensing reflectance [8–11]. The abundance of taxonomic groups of phytoplankton is another target of predictive models. The abundance of diatoms, prymnesiophytes (haptophytes), green algae and *Prochlorococcus* can be predicted using [Chl] [5]. Another model named PhytoDOAS uses remote sensing reflectance data at high spectral resolution for predicting the abundance of coccolithophores, dinoflagellates, cyanobacteria, and diatoms [12, 13]. Some models have been developed to predict the dominant size or taxonomic groups rather than the abundance [14–16]. One of this type of models named PHYSAT can predict communities dominated by diatoms, haptophytes, *Prochlorococcus* and *Synechococcus* defined by pigment concentration ratio [14, 15]. Another model has been developed to predict the distribution of the biogeochemical provinces [17]. However, zooplankton abundances have remained difficult to predict from satellite-derived data as they do not perform photosynthesis and tend to be transparent [18]. Nevertheless, some copepods harbor the carotenoid astaxanthin and their swarms can be observed from space [19].

These previous methods present a limitation regarding the numbers of defined plankton groups, as most of them are based on empirical relationships between pigments and light absorption. Even though these methods allow to provide a synoptic view on the spatiotemporal extent of the main groups of phytoplankton, they are lacking taxonomic resolution and do not reproduce the complexity of a planktonic community. In order to tackle this point, here, we present a machine learning trained model for global satellite observations of the representative communities as captured by a global ocean plankton network. Its targets are mixed auto-, hetero- and mixotrophic protist communities delineated from rDNA 18S V4 metabarcoding data at a high taxonomic resolution. Our approach is a network-oriented one, which was inspired by the Bayesian network model used to predict the metabarcoding-based bacterial composition in the English Channel [20]. There are two difficulties in predicting the species composition directly from satellite-derived data. The first difficulty is the substantial number of response variables as compared to predictor variables. There are hundreds of thousands of species represented in the metabarcoding dataset but only nine bands of visible light acquired by multispectral sensors are available as predictor variables. The second difficulty is the small number of samples. In this study, we used the largest available compilation of eukaryotic metabarcoding data, complemented with a novel sequence data from the *Tara* Oceans expeditions, but only a few hundred samples were available for analysis after appropriate filtration. By focusing on ecological networks, these two difficulties were alleviated by reducing the number of variables (dimensionality) in the metabarcoding data. Ecological networks tend to be structured, and are non-randomly assembled [21]. Indeed, a previous study showed through an unsupervised approach for community delineation that the global plankton network is self-organized by marine biomes [22]. We took advantage of this property of plankton networks to reduce dimensionality and convert the problem into a multiclass prediction.

## Materials and Methods

### Satellite data

Ocean color data acquired by the Moderate Resolution Imaging Spectroradiometer (MODIS) on board the Aqua and the Terra satellites were used in this study. Level-3 data, mapped to a 5□ (ca. 9 km on the Equator) square monthly grid, were downloaded from the Ocean Color Web operated by NASA (https://oceancolor.gsfc.nasa.gov/). The data included 17 parameters consisting of remote sensing reflectance (*R_rs_(*λ)) from 10 visible light wavelengths (412, 443, 469, 488, 531, 547, 555, 645, 667, and 678 nm), six environmental parameters derived from *R_rs_* ([Chl], diffuse attenuation coefficient for downwelling irradiance at 490 nm, particulate organic/inorganic carbon concentration, photosynthetically available radiation, and normalized fluorescence line height), and SST based on infrared measurements. The data were acquired from January 2003 to December 2021. To reduce the number of missing values, the data from both satellites were used. In case the values from both satellites were available for a grid, averaged values were used because they were well correlated (Fig. S1).

### Two-dimensional (2-D) projection of satellite-derived parameters

To capture the range of all possible satellite-derived parameter values, a 2-D projection was performed by randomly selecting grid cells. Twenty thousand grid cells were randomly selected from all the 5□ square grids. After removing grid cells on land or in coastal regions and those with missing data, 8,419 grid cells remained (Fig. S2). A sampling month was randomly selected from 120 months (January 2009 to December 2018) for each grid cell. The satellite-derived parameters for these randomly selected grid cells and months were standardized by subtracting the mean and scaling to unit variance. Finally, the 8,419 points with the 17 parameters were projected onto a 2-D map by Uniform Manifold Approximation and Projection (UMAP) using the Python package umap-learn [23].

### Metabarcoding data

Amplicon sequence data (837 127 965 million reads) targeting 18S V4 regions from 1 011 samples (1 191 datasets) collected through the *Tara* Oceans expeditions were generated and registered under the EMBL/EBI-ENA EukBank project. Raw sequencing data were downloaded from the EMBL/EBI-ENA EukBank umbrella project in their native format. When applicable, reads were merged and trimmed (vsearch [24], cutadapt [25]) to cover the 18S V4 region, as defined by the primers TAReuk454FWD1 and TAReukREV3, resulting in 347 327 830 unique sequences, representing 1 672 099 024 reads. After clustering (swarm [26], chimera detection (uchime [27]), quality-based filtering, and post-treatments based on occurrence patterns (swarm, lulu [28]; https://github.com/frederic-mahe/mumu), representative sequences were pairwise compared to the 18S rDNA database EukRibo [29], using a global pairwise alignment approach (usearch_global vsearch’s command), and taxonomically assigned to their best hit (https://github.com/frederic-mahe/stampa/). The filtered occurrence table contains 460 147 swarms (here after referred to as amplicon sequence variants (ASVs)), representing 1 403 019 176 reads, collected from 15 562 samples.

As for the usage of the EukBank occurrence table for the analysis, the raw number of reads was rarefied to 10 000 reads per sample. A total of 1 715 samples from the ocean surface (depth < 10 m) with spatiotemporal metadata were retained. These came from several ocean sampling projects such as *Tara* Oceans [1], Malaspina [30], and Australian Microbiome [31]. Occurrences in sequencing replicates from *Tara* Oceans were averaged. Samples from *Tara* Oceans were size fractioned into several classes (e.g. 0.8–5, 5-20, 20-180, and 180-2000 µm), but most samples from other projects were not size fractioned (0.2-3 µm or > 0.2 µm). The samples from four size fractions mainly targeting piconano-plankton (0.2– 3 µm, > 0.2 µm, 0.8–5 µm, and > 0.8 µm) were relatively similar in taxonomic composition (Fig. S3) and were used in this study to maximize the number of samples available for analysis. They were averaged inside each of the 653 bins that match one-by-one with the 5□ square monthly satellite data grid. Although more than one sample from different size fractions, sampling locations and times were assigned to a single bin, samples in a same bin were more similar compared to samples from different bins (Fig. S4). Hereafter, we call these bins “samples”.

### Spatial resampling

A total of 653 metabarcoding samples from previous processing were further filtered using the following procedure. Samples with missing satellite data values owing to bad weather or other reasons were removed. Samples from locations where the sea floor was shallower than 200 m were classified as coastal samples and were removed following previous recommendations [32] using a global relief model [33]. Samples were thinned so that they were separated by a minimum of 200 km from each other, using the R package spThin [34]. This procedure resulted in 177 samples available for analysis (Fig. S5).

### Network inference

ASVs were selected by their occurrence to reduce the number of ASVs to those similar to previous studies that analyzed network structures [35, 36]. Two hundreds and eight ASVs with a minimum occurrence larger than 0.2% (20 reads) in at least 10% of samples (18 samples) were retained (Figs. S6). ASV read counts were centered log-ratio (clr) transformed [37]. An ecological network was inferred based on co-occurrence patterns using the Julia package FlashWeave [38] with the settings “heterogeneous=False”, “sensitive=True”, and “alpha=0.05” as in previous studies [38, 39]. FlashWeave is a package for calculating partial Pearson’s correlation coefficient between ASV pairs using a recursive approach. The nodes in the obtained network were ASVs, and the edges were made based on correlations between ASV pairs. Only positive correlations (edges) were considered here. For network community detection in the network, Fast Greedy, Infomap, Label Propagation, Leading Eigenvector, Leiden, Louvain, Spinglass, and Walktrap algorithms were applied using the R package “igraph” (https://igraph.org/). To measure the structure of the detected community division, we used the modularity index Q as defined by the following equation:

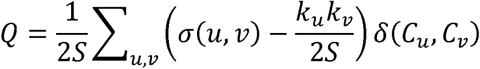

where *u*, *v* are nodes (ASVs), *σ*(*u,v*) is an edge weight (partial correlation coefficient) between *u* and *v*, *S* is the sum of all edge weights, *k_u_* is a weighted degree of node *u*, *C_u_* is a community to which node *u* belongs, and *δ*(*x,y*) is 1 if *x*=*y* and 0 otherwise [40].

### Edge satisfaction

We defined an “edge satisfaction index” to determine which community dominated each sample. If *C* is a community and *i* is a sample, then the edge satisfaction index of *C* and *i* is defined by

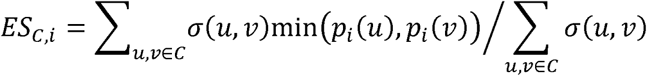

where *u*, *v* are nodes, *σ*(*u,v*) is an edge weight between *u* and *v*, *P_i_*(*u*) is a weight of node *u*, which is the sigmoid transformation of the clr-transformed read count of ASV *u* in sample *i*. Briefly, this index measures the ratio of the number of edges between existing nodes in a given sample and the number of all the edges within a given community. The nodes and edges have a weight between 0 to 1 (because only positive correlations were considered). The edge satisfaction index is thus also between 0 to 1.

### Machine learning and cross-validation

Several machine learning algorithms were used to train predictive models of the representative community based on satellite-derived data. In addition to the satellite-derived parameters, spatial parameters (longitude and latitude) were also tested for their ability in the prediction. The sine and cosine of the longitude were used as independent parameters because it is circular (−180° and 180° are the same). K-nearest Neighbors, Naïve Bayes, Multilayer Perceptron, Random Forest, and Support Vector Machine (SVM) were applied using the Python package “scikit-learn” (https://scikit-learn.org/). In the training process for all the methods except Random Forest, the satellite-derived and spatial parameters were standardized by subtracting the mean and scaling to unit variance. The hyperparameters that were tuned with the grid search are shown in Table S1. Both leave-one-out cross-validation and buffered cross-validation [41] were used to measure the model accuracy. In the buffered cross-validation, a test sample is chosen like leave-one-out, but samples inside a buffer region among the test sample were excluded from training samples. The buffer was set to 2,000 km radius. In each fold of the training, hyperparameters were chosen with exhaustive search using the implementation of grid search in scikit-learn. The class prediction output of each method was used to measure accuracy, and output probabilities were used to calculate the receiver operating characteristic (ROC) curve.

### Time series prediction

The predictive model was trained again with all 177 samples. A 5-fold grid search was used to choose hyperparameters. To reduce the computational cost, a grid cell at the center of each 12 by 12 grids square was chosen. In other words, a grid cell was chosen per every 1° square monthly grid cells because the original grid is 5□ square. The trained model was applied to this 1° square grid data set for January 2003 to December 2021.

## Results

### 2-D map of points with 17 satellite-derived parameters

We generated a 2-D map of points with 17 satellite-derived parameters using UMAP to observe the parameter ranges (Fig. 1). More than eight thousand points were used to train UMAP. These points were randomly selected from all possible locations and times to document the shape of the “continents” in UMAP, which represent the possible range of values of the satellite parameters (Fig. S2). Points associated with the EukBank metabarcoding samples were scattered among most of the regions in the UMAP continents (excluding a region indicated by an arrow). This result indicates that the metabarcoding data covered a wide range of the parameter space and are suitable for being analyzed in terms of their relationship with satellite data, although the number of samples was not large.

**Figure 1.**
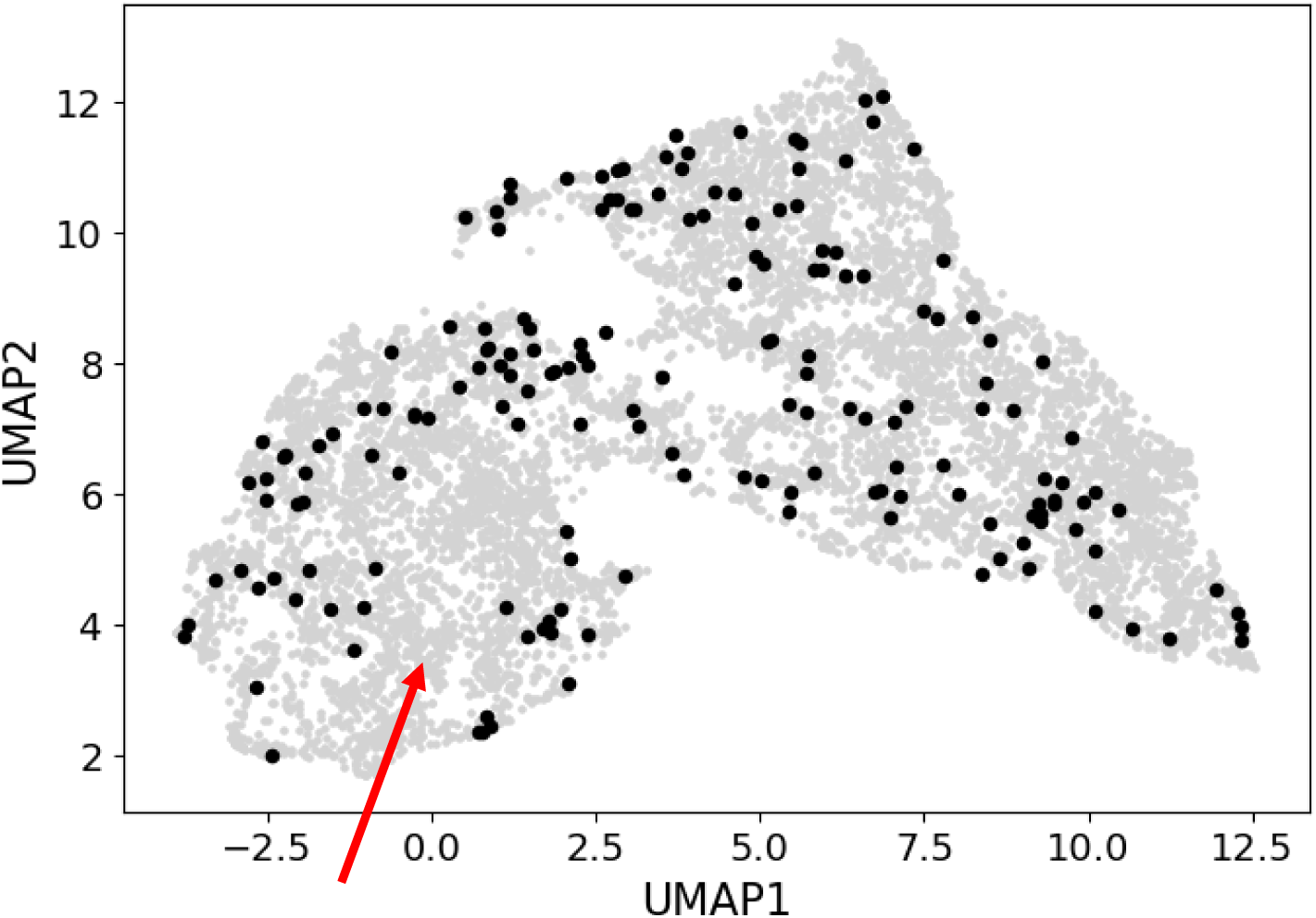
Metabarcoding samples used for prediction model training. Points associated with metabarcoding samples projected on the 2-D map of the satellite-derived parameter space (black points). Grey points are randomly selected grid cells used for learning the map. Red arrow indicates the region without metabarcoding samples.

### Network inference and community detection

The ecological network based on ASV co-occurrence patterns was inferred using the FlashWeave algorithm. In the network, 560 positive edges (correlation coefficients > 0) between 208 ASVs were detected (Fig. 2A). We applied several community detection algorithms on the network. The communities detected by Leiden and Spinglass algorithms had the highest modularity index (0.55) (Fig. S7). In the following analysis, the communities detected by the Leiden algorithm [42] were used because it captured the macro structure better than the others (i.e., there were no small communities) (Fig. S7). Among the six detected communities, community 1 was well separated from the other five communities, which formed one super community having highly aggregated community structure (Fig. 2B). In the super community, communities 2 and 5 were strongly connected with communities 3 and 6, respectively (Fig. 2B).

**Figure 2.**
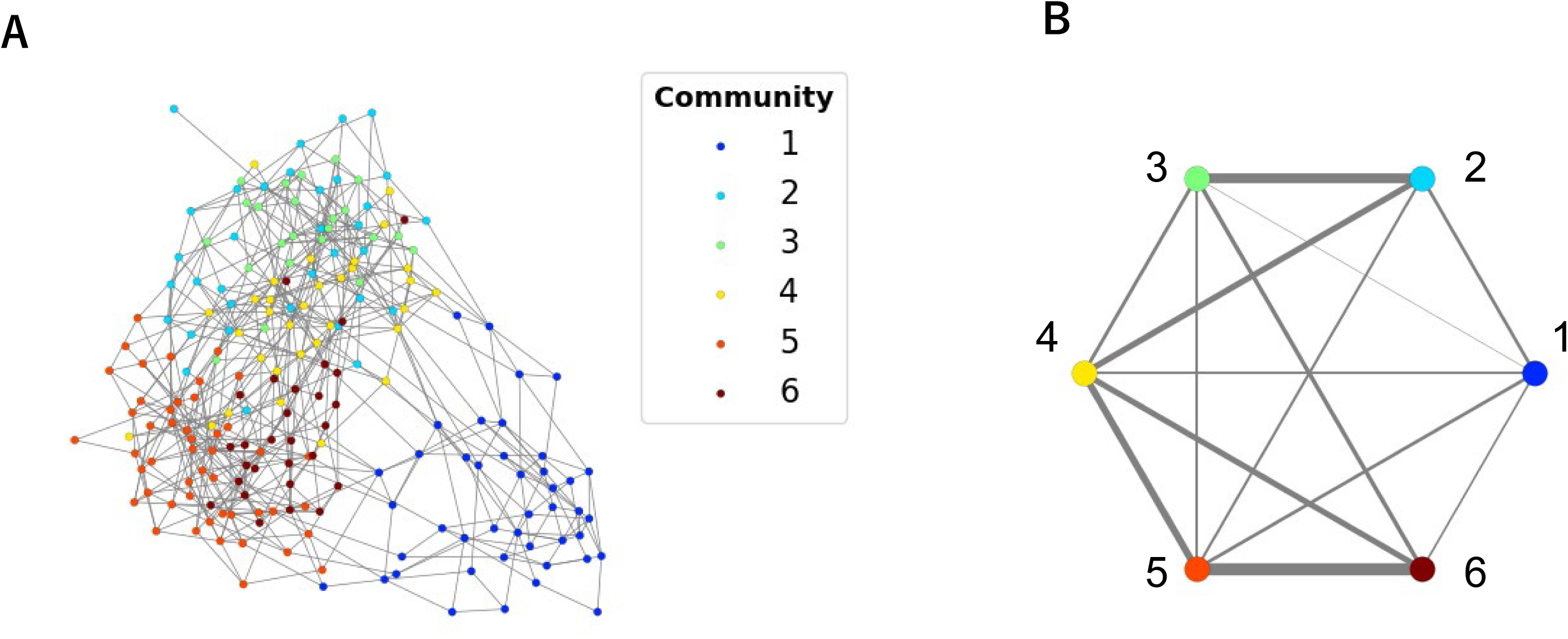
Plankton network inferred using metabarcoding data. **A** A force-directed representation of the network. Nodes (plankton ASVs) are colored by belonging community. **B** A graph representing community connection of same network. Width of edges are proportion to the number of inter-community edges.

The taxonomic breakdown of each community is shown in Table 1 and Fig. S8. The well-separated community 1 mainly consisted of Dinoflagellata (mainly Dinophyceae), but Dictyochophyceae (silicoflagellates) and Prymnesiophyceae (haptophytes) were also included. Other five communities had different characteristics in terms of taxonomy. Communities 5 and 6 were dominated by Dinoflagellata (mainly MALV-I and MALV-II), but communities 2 and 3 contained some Arthropoda. Community 4 consisted of half Dinoflagellata and half a variety of other taxa. See Data S1 for taxonomic annotation and assigned community for each ASVs.

**Table 1.** Taxonomic breakdown of plankton communities.

| Taxogroup 1<sup>a</sup> | Taxogroup 2<sup>a</sup> | 1 | 2 | 3 | 4 | 5 | 6 | Total |
| --- | --- | --- | --- | --- | --- | --- | --- | --- |
| Dinoflagellata | Dinophyceae | 16 | 16 | 3 | 3 | 6 |  | 44 |
|  | MALV-I | 4 | 1 | 6 | 7 | 19 | 6 | 43 |
|  | MALV-II | 5 | 1 |  | 6 | 15 | 11 | 38 |
|  | MALV-III | 2 |  | 4 |  | 1 |  | 7 |
|  | MALV-V |  |  |  |  | 1 | 3 | 4 |
| Metazoa | Arthropoda | 1 | 10 | 3 |  |  |  | 14 |
|  | Cnidaria |  | 3 |  |  |  |  | 3 |
|  | Urochordata |  |  | 1 |  |  |  | 1 |
| Ochrophyta | Dictyochophyceae | 2 |  |  | 1 |  | 1 | 4 |
|  | Pelagophyceae | 1 |  |  |  |  |  | 1 |
|  | MOCH-2 |  |  |  | 1 |  |  | 1 |
|  | Diatomeae |  |  | 1 |  |  |  | 1 |
| Chlorophyta | Mamiellophyceae | 1 |  |  | 3 |  |  | 4 |
|  | Chloropicophyceae |  |  | 2 |  |  |  | 2 |
| Prymnesiophyceae | Prymnesiophyceae | 3 | 1 |  | 1 |  |  | 5 |
| Radiolaria | Acantharea |  | 2 |  |  |  |  | 2 |
|  | RAD-B-clade |  |  | 1 | 1 |  |  | 2 |
|  | Spumellaria |  |  |  |  |  | 1 | 1 |
| MAST-01 | MAST-01 | 1 |  |  | 2 |  | 2 | 5 |
| Picozoa | Picozoa | 2 |  |  | 1 | 2 |  | 5 |
| Sagenista | MAST-04 |  |  |  | 2 |  | 2 | 4 |
|  | MAST-07 |  |  |  |  |  | 1 | 1 |
| Cryptomonadales | Cryptomonadales | 2 |  |  | 1 |  |  | 3 |
| Opalozoa | Nanomonadea |  |  |  | 1 |  | 1 | 2 |
|  | Bicosoecida |  |  |  |  | 1 |  | 1 |
| core-kathablepharids | core-kathablepharids | 1 |  | 1 |  |  |  | 2 |
| Ciliophora | Spirotrichea | 1 |  |  |  |  |  | 1 |
| Centroplasthelida | Centroplasthelida |  |  |  |  | 1 |  | 1 |
| Unclassified | Unclassified | 1 |  | 1 | 2 | 2 |  | 6 |
| Total |  | 43 | 34 | 23 | 32 | 48 | 28 | 208 |

a: Taxonomic level in the EukRibo.

### Representative community of samples

The newly proposed edge satisfaction index was used to measure the completeness of the communities in each sample (see methods). Briefly, this index is one in case all edges within a given community exist, and it is zero in case no edges exist. Fig. 3A shows the edge satisfaction index of all the samples and communities. Notably, community 1 was an exclusively assigned community for some of the samples. The community with the highest edge satisfaction index was considered as the representative community of the sample. The geographic distribution of the representative communities is shown in Fig. 3B. Community 1 was associated with high latitude regions, including the Arctic and the Southern Oceans. Communities 3 and 6 were mainly seen in tropical regions of the Pacific and the Indian Oceans, respectively. The other three communities were associated with mid-latitude regions.

**Figure 3.**
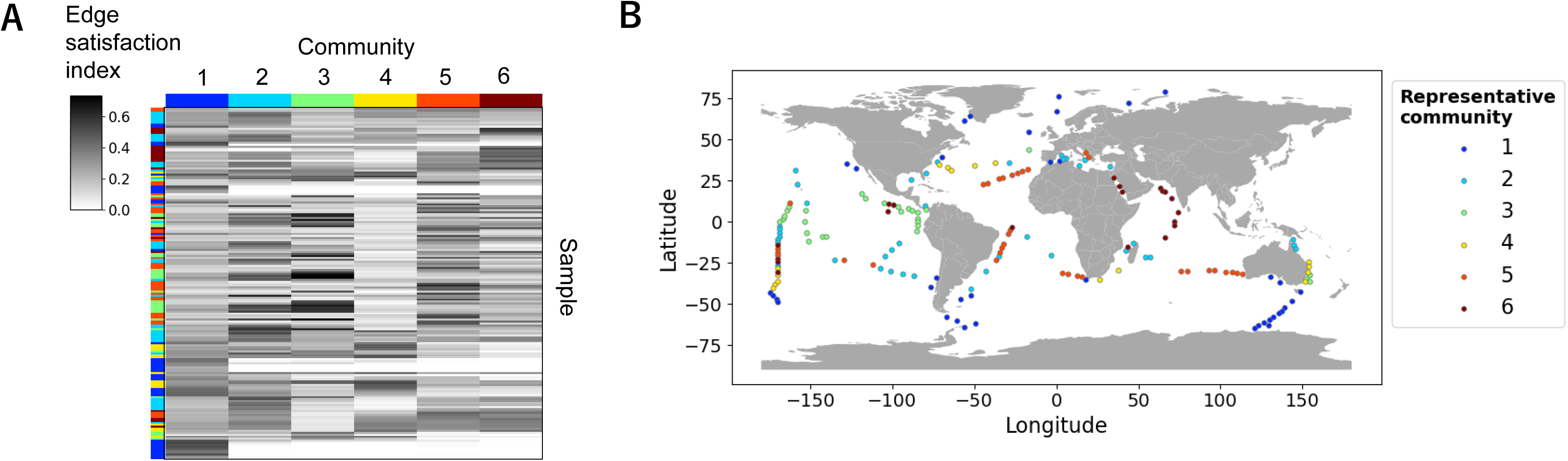
Representative community of samples. **A** A heatmap of edge satisfaction index. The left most column shows representative community of each sample. **B** Geographic distribution of representative communities.

In the 2-D map of satellite parameter space, samples formed clusters of representative communities (Fig. S9). For example, communities 1 and 5 dominated the bottom of the small and large continents of the map, respectively. This distribution implies a relationship between the satellite parameters and the representative communities.

### Prediction performance

We applied several machine learning algorithms to classify the representative communities based on satellite parameters. Among five machine learning methods we used, SVM achieved the highest prediction accuracy and micro-average area under the ROC curve (ROC-AUC) (Table S2). Using leave-one-out cross-validation, the accuracy and the ROC-AUC of SVM were 0.67 and 0.90, respectively (Figs. 4A and 4B). Using buffered cross-validation, which excluded neighbors of a test sample from training samples, the measures were reduced to 0.54 and 0.83, respectively (Figs. 4C and 4D).

**Figure 4.**
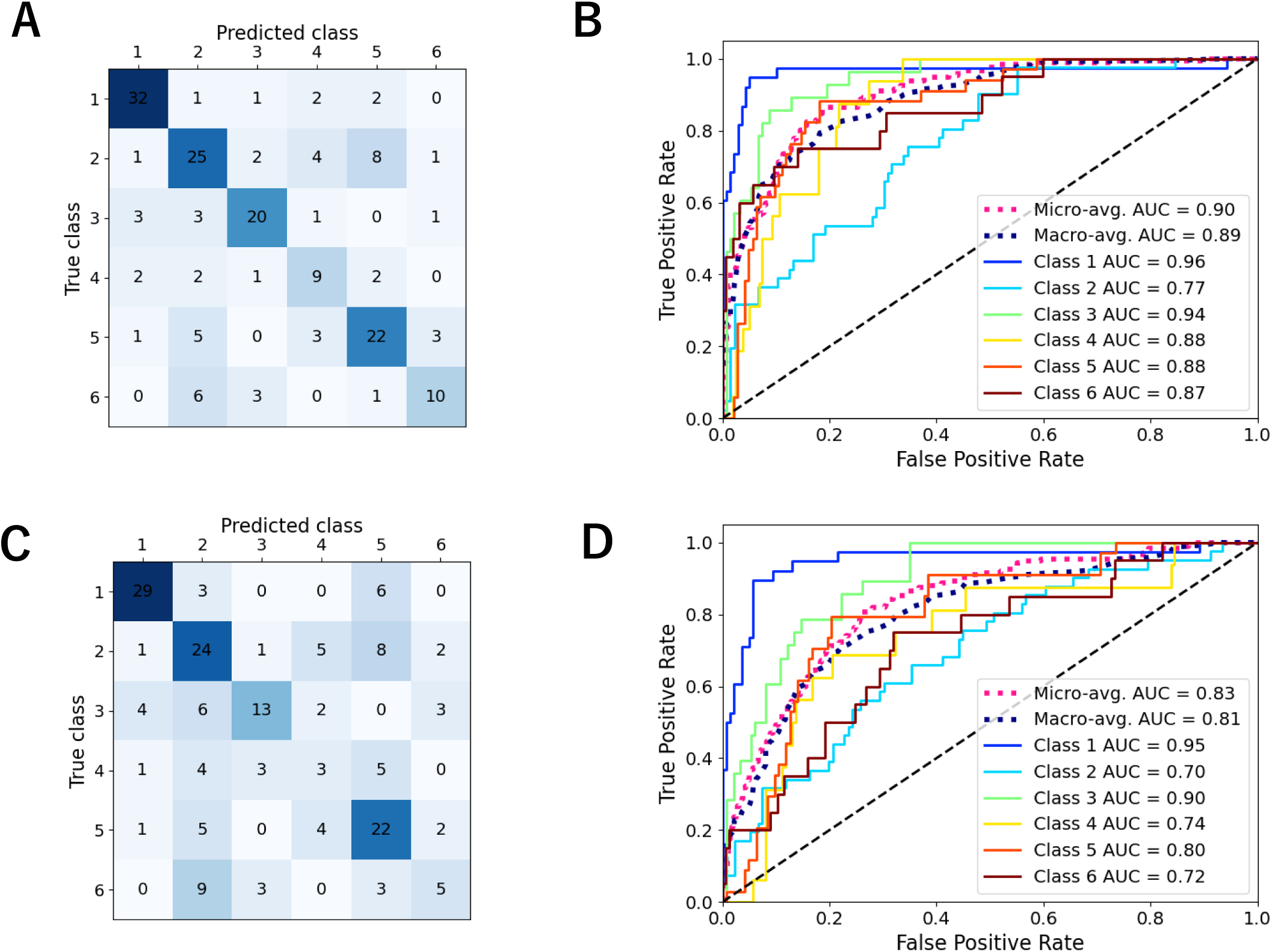
Performance of support vector machine on community prediction based on satellite-derived parameters. Performance of SVM using all the satellite-derived parameters. **A-B** The confusion matrix (**A**) and the ROC curve (**B**) in the condition of leave-one-out cross-validation. **C-D** The confusion matrix (**C**) and the ROC curve (**D**) in the condition of buffered cross-validation.

We compared the prediction performance when different sets of satellite-derived and spatial parameters were used (Table 2, Figs S10 and S11). For the prediction only using spatial parameters (latitude and sine/cosine of longitude), the ROC-AUC dropped from 0.91 to 0.59 (close to 0.50, i.e., random prediction) when we changed the cross-validation method from leave-one-out to buffered. In contrast, there was a small decrease from 0.90 to 0.83 for the prediction with the satellite-derived parameters. This result indicates the advantage of using satellite-derived parameters to classify the representative communities when spatial biases are appropriately controlled. The prediction performance with only SST or [Chl] was not as good compared to the one with all satellite-derived parameters, but it was improved when SST and [Chl] were combined. Adding other five environmental parameters to SST and [Chl] further improved the performance but it was still slightly worse than that with all satellite-derived parameters (including SST, environmental parameters and *R_rs_*).

**Table 2.** Comparison of prediction performance using different sets of satellite-derived and spatial parameters. Support vector machine were used in all the parameter sets.

| Parameter set | Leave-one-out cross-validation |  | Buffered cross-validation |  |
| --- | --- | --- | --- | --- |
|  | Accuracy | ROC-AUC<sup>a</sup> | Accuracy | ROC-AUC<sup>a</sup> |
| All satellite-derived parameters | 0.67 | 0.90 | 0.54 | 0.83 |
| Latitude, Longitude<sup>b</sup> | 0.68 | 0.91 | 0.29 | 0.59 |
| SST | 0.40 | 0.79 | 0.28 | 0.72 |
| [Chl] | 0.43 | 0.72 | 0.23 | 0.62 |
| SST, [Chl] | 0.52 | 0.86 | 0.47 | 0.82 |
| SST, Environmental parameters | 0.58 | 0.88 | 0.50 | 0.83 |

a: Micro-average area under the ROC curve.
b: Sine and cosine of longitude were used as parameters.

### Time series prediction

A full SVM model of community prediction was trained with all 177 samples. A 5-fold grid search selected the linear kernel and the L2 penalty parameter C=1.0 for the full model. The SST was the most important parameter in the full model, followed by photosynthetically available radiation, and several channels of *R_rs_* (Fig. S12). The chosen threshold of the probabilistic output of SVM was 0.28, which gave the highest F1 score in cross-validation (Fig. S13).

We applied the full SVM model to predict a 19-year time series, from January 2003 to December 2021, of community distributions (Movie S1). The global community distributions in each season of year 2021 are shown in Fig. 5. Communities 1 and 5 are located at high- and mid-latitudes, respectively. Communities 3 and 6 are tropical. Community 2 fills the gap between communities 5 and 3. Community 4 shows the pattern related to warm currents. It is related to the region of the extensions of the Kuroshio and Gulf Stream in the late autumn and early winter of the northern hemisphere (November–January) and that of the Brazil, Agulhas, and East Australian Currents in the late autumn and early winter of the southern hemisphere (May–July).

**Figure 5.**
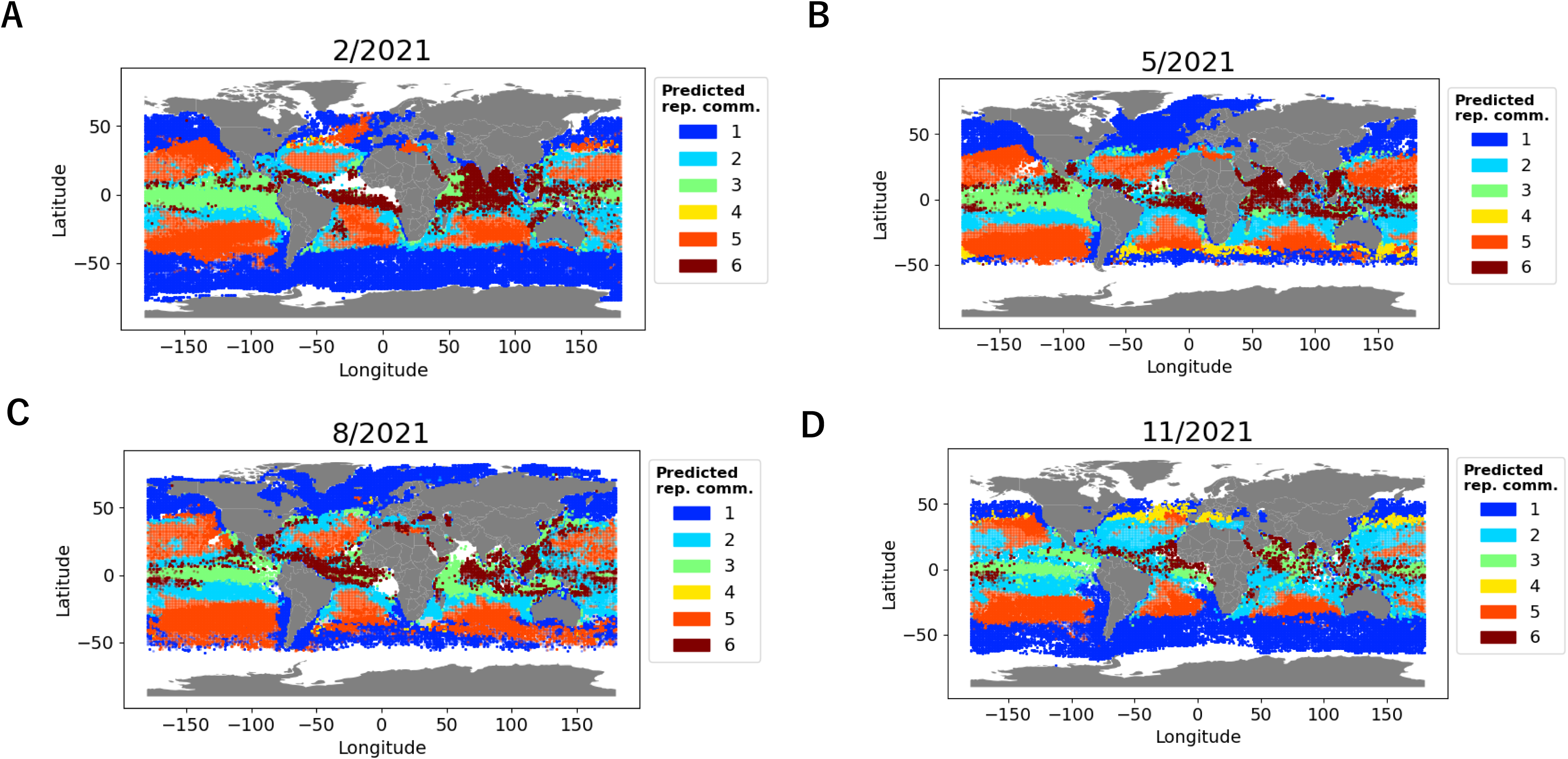
Spatiotemporal distribution of communities predicted from satellite-derived parameters. Representative community distribution in February (**A**), May (**B**), August (**C**), and November (**D**) of year 2021 predicted from satellite-derived parameters. When multiple communities were predicted to be representative in the same point, community with the highest probability is show in pale color. Grey point means no community was predicted to be representative.

## Discussion

Here, representative plankton community networks inferred from rDNA 18S V4 global metabarcoding data from EukBank were successfully predicted from satellite data using a machine learning approach. The outputs of this model are the plankton communities in a way similar to the taxonomic group output of the PHYSAT model [15] rather than a quantitative abundance output like the PhytoDOAS model [12, 13]. Our method has two advantages over these latter models. First, the representative community output from our method included all the taxonomic range of microbial eukaryotes from phytoplankton and zooplankton to heterotrophic protists. There have been some attempts to capture zooplankton abundance from satellite data [19, 43], but it is still difficult to capture a global view over the wide taxonomic range. Our network approach can be extended to prokaryotes and viruses (as they are strongly associated with eukaryotic communities [44, 45]), which are also difficult to observe from satellites due to their small size and lack of optical properties.

Second, the output of our model was directly connected with the ASVs inferred from the metabarcoding data. An ASV is a unit with a high taxonomic resolution that is operationally treated as a “species”; thus, the representative communities integrate high taxonomic resolution information. For example, dinoflagellates were treated as one group in the PhytoDOAS model [12], whereas they were represented by 136 ASVs that were classified into one of the six different communities in this study (Table 1, Fig. S8 and Data S1). Although the high taxonomic resolution of metabarcoding data is fascinating, handling their limited number of samples and high dimensionality lead several limitations. We used only four size fractions mainly targeting piconano-plankton (0.2–3 µm, > 0.2 µm, 0.8–5 µm, and > 0.8 µm) to maximize the number of samples available for analysis. By this procedure, taxonomies only observed in larger size fractions (e.g. diatoms) were not included in the network (Fig. S3A). We choose the criteria of ASV selection and the resolution of community detection (e.g. resolution parameter of Leiden algorithm) to reduce the high dimensionality of the metabarcoding data. This choice of parameters naturally reduced the precision and true diversity of detected communities. A benchmarking test of community detection will overcome this limitation in the future.

Our results indicated that the performance of the prediction based on SST and [Chl] was relatively high (Table 2, Figs. S10 and S11). This is not unexpected as SST and [Chl] correlate with microbial community structure in the ocean [1]. We also showed that the prediction performance with all 17 satellite-derived parameters (including SST, environmental parameters and *R_rs_*) was higher than with only on SST and/or [Chl] (Table 2, Figs. S10 and S11). This result suggested the advantage of using additional environmental parameters and *R_rs_* to predict communities, but the improvement of the performance was not large. Hyperspectral *R_rs_* from future global satellite missions such as PACE [46] will likely improve prediction performance.

We used 177 samples (after an appropriate filtration), which was relatively small for applying a machine learning approach. This may explain why the linear SVM was the best predicting algorithm for our problem. More complex and nonlinear algorithms such as Multilayer Perceptron, Random Forest, and kernel SVM overfitted the training dataset during full model training (Fig. S14). Although it is not easy to increase the number of samples because of cost, labor, and weather limitations for sampling cruises, the current number, 1,715 (and only 177 after binning and thinning) out of 10,772 samples contained in the metabarcoding dataset is quite limited. Thus, plankton samples from the ocean surface should be strategically collected in the future. The 2-D map of satellite-derived parameters for the ocean (Fig. 1) can provide guidance for such strategical guided sampling. If there is a one-to-one connection between the satellite data and the microbial community assemblies, regions without EukBank samples in the map will relate to communities that have yet to be observed and will be an important target for future sampling. A region on the map indicated by an arrow is an example of this kind of unexplored region (Fig. 1). We found that this region is mainly consisted of points associated with high latitudinal regions of the southern hemisphere and the North Pacific Ocean (Fig. S15).

The time series prediction of communities using the trained model revealed the spatiotemporal distribution of each community (Fig. 5 and Video S1). Generally, community distributions had a rough correspondence with Longhurst biomes [47] (Fig. S16). Community 1 corresponded with the “polar” biome, community 5 corresponded with the “westerlies” biome, and communities 3 and 6 corresponded with the “trades” biome. The same distribution can be seen in the representative community of each sample used to train the model (Fig. 3B). Considering the latitudinal self-organization previously observed and described in plankton community networks [22], this correspondence showed that the newly proposed edge satisfaction index could appropriately capture the representative communities. Community 4 had a seasonal spatiotemporal distribution possibly related to the extensions of the western boundary currents (Fig. 5 and Video S1). A previous study showed that seasonal changes in environmental variables (phosphate, nitrate, silicate, and dissolved inorganic carbon) were the highest in the extension of the Kuroshio among other regions in the Pacific basin [48]. Furthermore, clear seasonal variations in the abundance of cyanobacterial diazotrophs were observed in the same region [49]. Community 4 connected the two well-connected pairs (communities 2 and 3, 5 and 6) of the super community in the network (Fig. 2B) and had relatively high taxonomic diversity (Table 1 and Fig. S8). In a simulation of emergent phytoplankton in the ocean, areas downstream of the western boundary currents showed high species diversity [50]. The target of our model is the global spatiotemporal distribution of communities, while our network-based approach will be applicable to satellite observations of local ecosystems (e.g., Hawaii and Bermuda).

In this study, we inferred the ecological network of ASVs using a global metabarcoding dataset and identified six distinct communities. We applied SVM to train the predictive model of these communities based on satellite data and obtained an accuracy of 67%. The spatiotemporal distribution of these communities was shown by applying the model to 19 years of global satellite data. Our model was able to predict communities that included phytoplankton, zooplankton, and heterotrophic protists. The network-oriented approach used in this study can be easily extended to identify the distribution of prokaryotes and viruses. Given the ability of the model to predict the spatiotemporal dynamics of plankton communities from space, our combined network-based and machine learning approach provides a particularly useful tool to monitor and survey the impact of environmental and climate change on plankton communities at both local and global scale.

## Data Availability Statement

Newly sequenced *Tara* Oceans 18S V4 data have been deposited to EMBL/EBI-ENA: PRJEB6610 (*Tara* Oceans), PRJEB9737 (TARA Oceans Polar Circle).

## Acknowledgments

We thank the *Tara* Oceans consortium, the EukBank consortium, and the people and sponsors who supported the *Tara* Oceans Expedition (http://www.embl.de/tara-oceans/) for making the data accessible. This is contribution number XXX of the *Tara* Oceans Expedition 2009–2013. Computational time was provided by the Supercomputer System, Institute for Chemical Research, Kyoto University. This work was supported by JSPS/KAKENHI (Nos. 18H02279 and 19H05667 to H.O.), and the Collaborative Research Program of the Institute for Chemical Research, Kyoto University (2020-29 to K.T.). We thank Leonie Seabrook, PhD, from Edanz (https://jp.edanz.com/ac) for editing a draft of this manuscript.

## Author Contributions

H.K. designed the study, performed most of the bioinformatics analyses and wrote the initial manuscript. H.E., R.N., K.T., and H.O. contributed to the design of the work and supervised H.K. N.H., C.B., F.M., and C.d.V. performed the amplicon sequence data processing and annotation. J.P., K.L., O.B., and P.W. treated biological samples and performed sequencing. R.E.H., S.C., L.K.-B., E.B., and C.B. provided expertise in marine biology. *Tara* Oceans Coordinators (S.G.A., M.B., P.B., E.B., C.B., G.C., C.d.V., G.G., L.G., N.G., P.H., D.I., O.J., S.K., L.K.-B., E.K., F.N., H.O., N.P., S.P., C.S., S.S., L.S., M.B.S., S.S., and P.W.) contributed to expeditionary infrastructure needed for global ocean sampling, sample processing, and data production. All authors contributed to the interpretation of data and finalization of the manuscript.

---

**Table S1.** Machine learning methods and their hyperparameters.

| Method | Scikit-learn implementation | Hyperparameters |
| --- | --- | --- |
| K-nearest Neighbors | sklearn.neighbors.KNeighborsClassifier | Number of neighbors (1, 2, ..., 9) |
| Naïve Bayes (Gaussian) | sklearn.naive_bayes.GaussianNB | Smoothing parameter ($10^{-4}$, $10^{-3}$, ..., $10^3$) |
| Multilayer Perceptron (one hidden layer) | sklearn.neural_network.MLPClassifier | Activation function (identity, logistic, tanh, ReLU), Hidden layer size (10, 20, 30, 40), L2 penalty parameter ($10^{-6}$, $10^{-5}$, ..., $10^3$) |
| Random Forest | sklearn.ensemble.RandomForestClassifier | Number of trees (10, 100, 1000, 10000) |
| Support Vector Machine | sklearn.svm.SVC | Kernel (linear, RBF), Coefficient of RBF kernel ($10^{-4}$, $10^{-3}$, ..., $10^3$), L2 penalty parameter ($10^{-4}$, $10^{-3}$, ..., $10^3$) |

**Table S2.** Comparison of machine learning methods. All the satellite-derived parameters were used in the validations.

| Method | Leave-one-out cross-validation |  | Buffered cross-validation |  |
| --- | --- | --- | --- | --- |
|  | Accuracy | ROC-AUC<sup>a</sup> | Accuracy | ROC-AUC<sup>a</sup> |
| K-nearest Neighbors | 0.56 | 0.86 | 0.47 | 0.76 |
| Naïve Bayes | 0.55 | 0.85 | 0.50 | 0.81 |
| Multilayer Perceptron | 0.60 | 0.88 | 0.54 | 0.82 |
| Random Forest | 0.63 | 0.89 | 0.50 | 0.81 |
| Support Vector Machine | 0.67 | 0.90 | 0.54 | 0.83 |

a: Micro-average area under the ROC curve.

**Figure S1.**
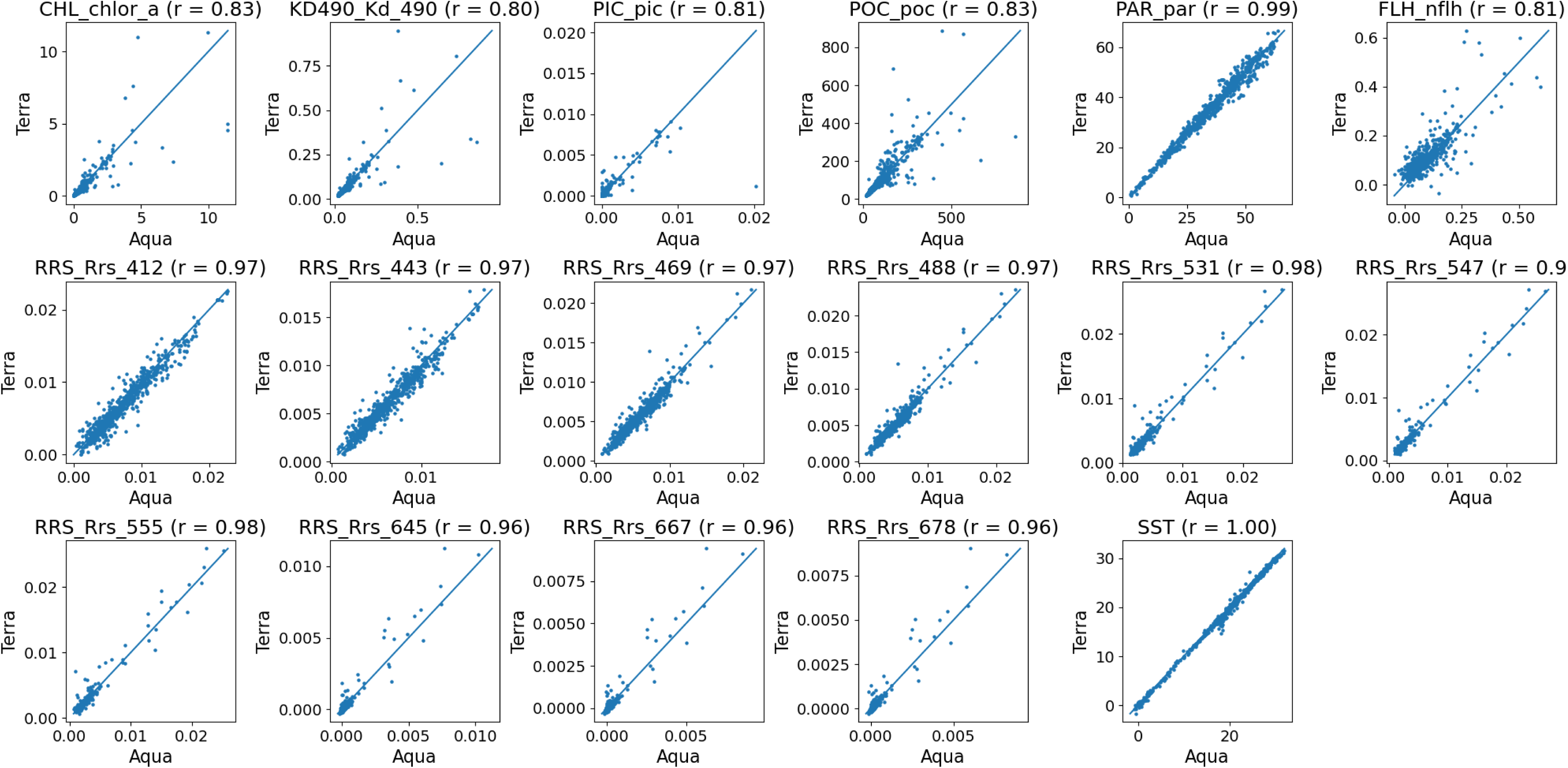
Comparison of satellite-derived parameters acquired by the Aqua and the Terra satellite. Pearson’s correlation coefficient was shown for each parameter pairs.

**Figure S2.**
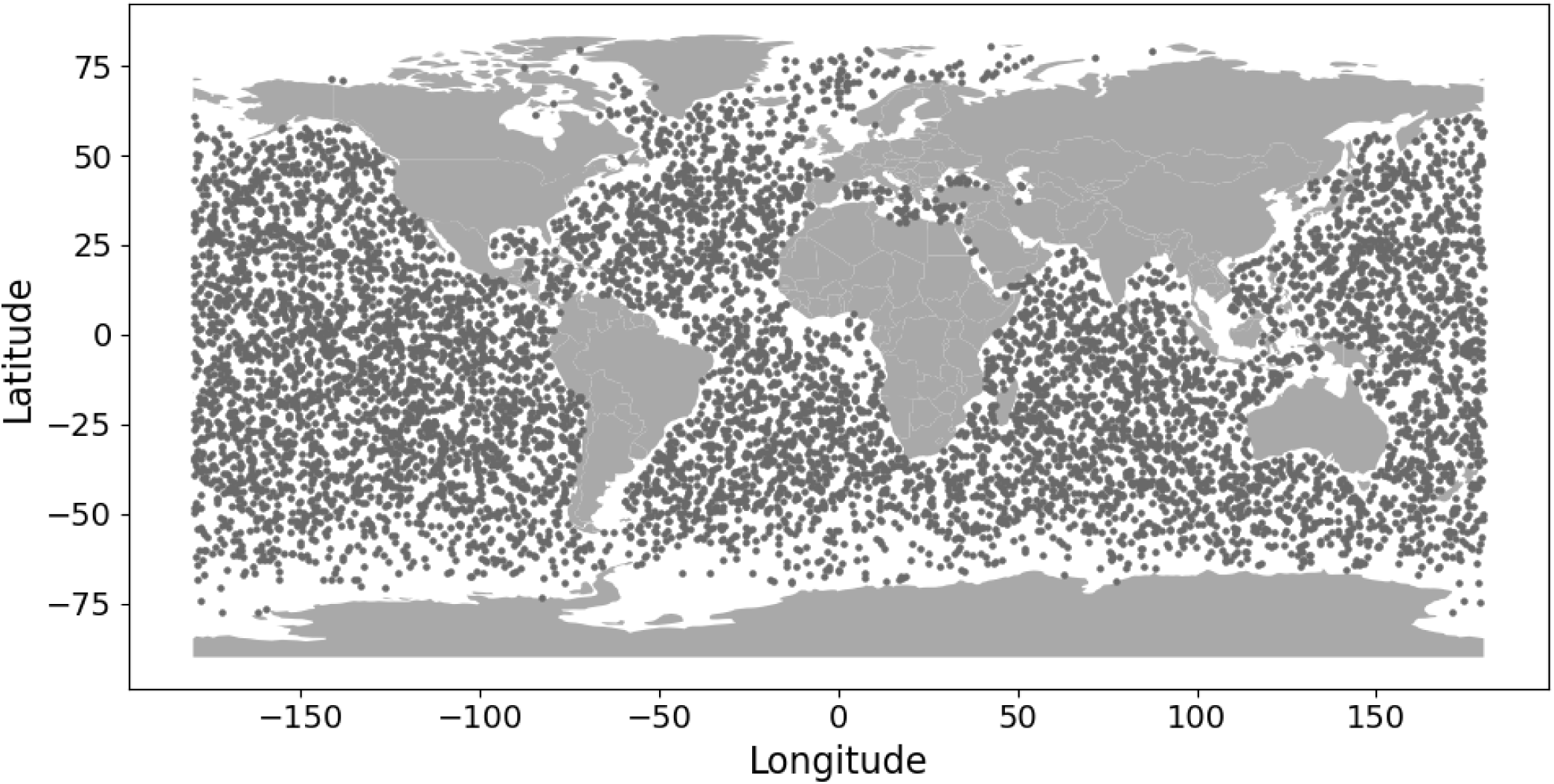
Location of satellite-derived parameter samples used for leaning UMAP. Dark grey points are location of grid cells used to learn a map with UMAP. Sampling month was also randomly selected for each grid cell.

**Figure S3.**
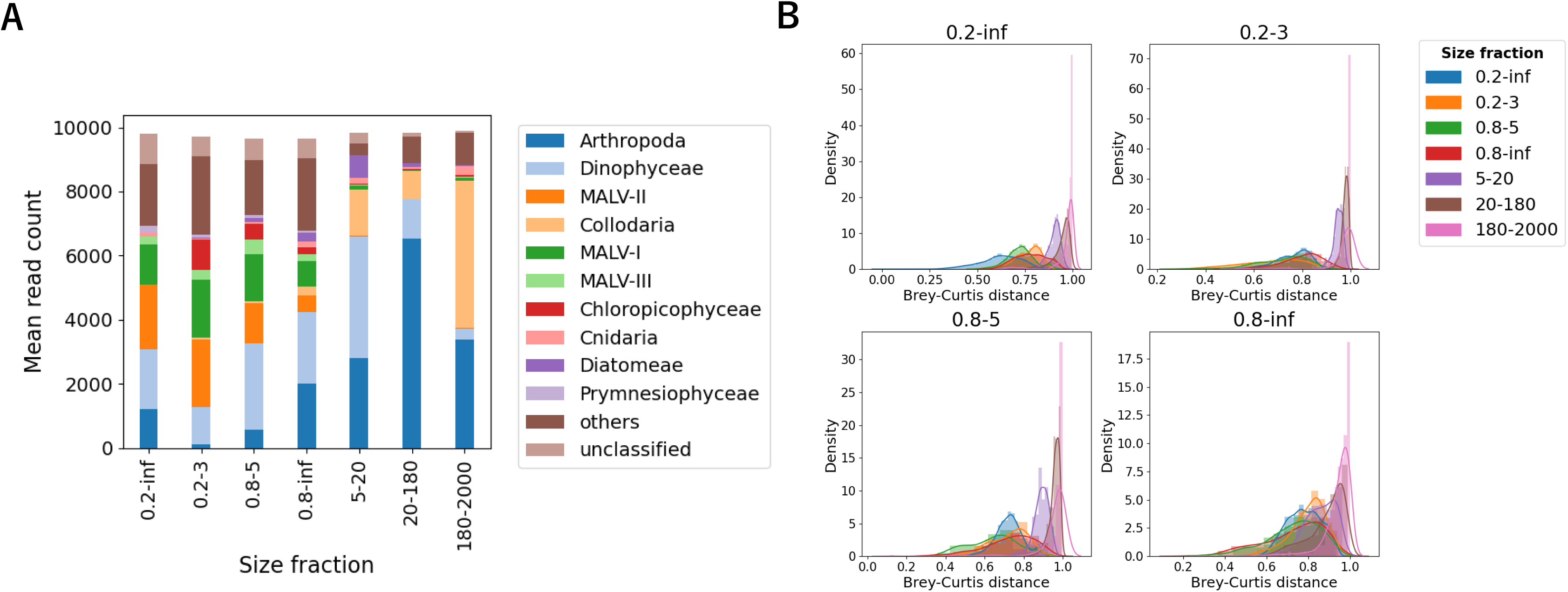
Comparison of size fractions in the South Pacific Subtropical Gyre. Taxonomically difference of size fractions was checked among samples from the South Pacific Subtropical Gyre, which contains samples from all major size fractions. **A** Mean **t**axonomic composition of each size fraction. Taxonomic level is “taxogroup 2” in the EukRibo. **B** Comparison of taxonomic distance of intra- and inter-size-fraction samples. Brey-Curtis distance was calculated based on ASV read count.

**Figure S4.**
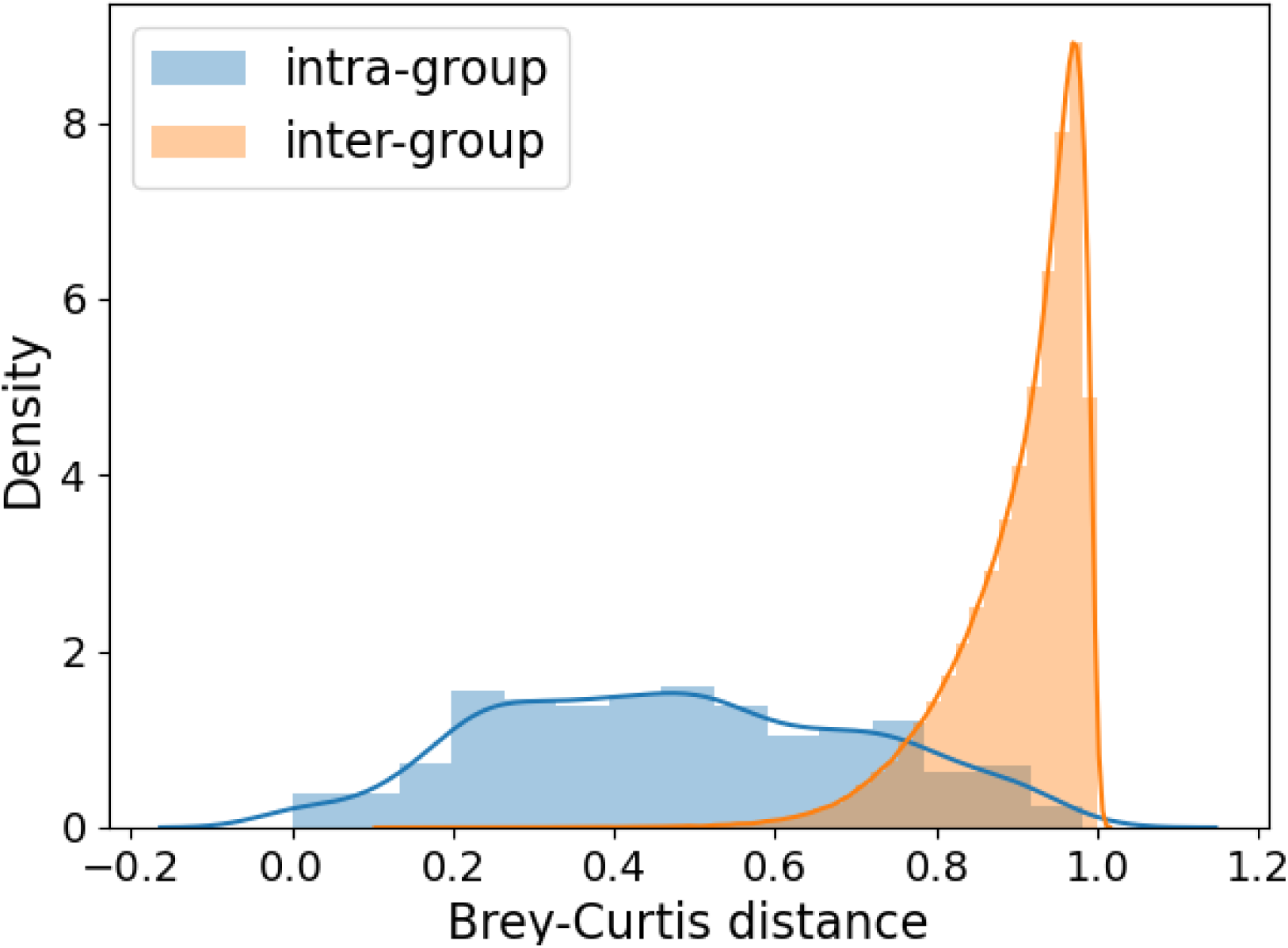
Comparison of intra- and inter-bin sample distance. Brey-Curtis distance was calculated based on ASV read count. Intra-bin sample distance is small enough compared to inter-bin sample distance in the most cases.

**Figure S5.**
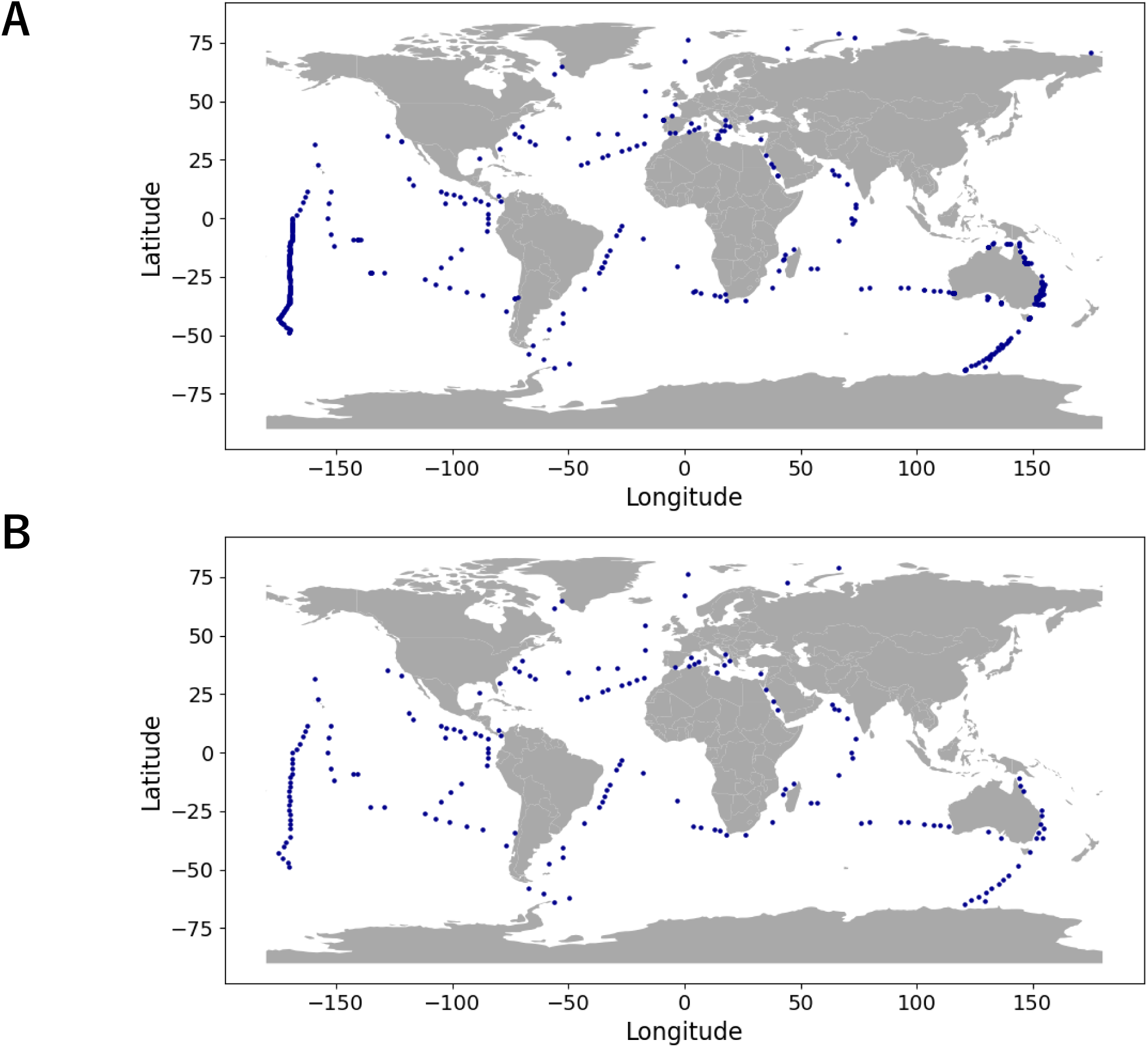
Spatial resampling of metabarcoding data. **A** Geographic location of 653 metabarcoding samples (bins) before spatial resampling. **B** 177 samples retained and used for the analysis.

**Figure S6.**
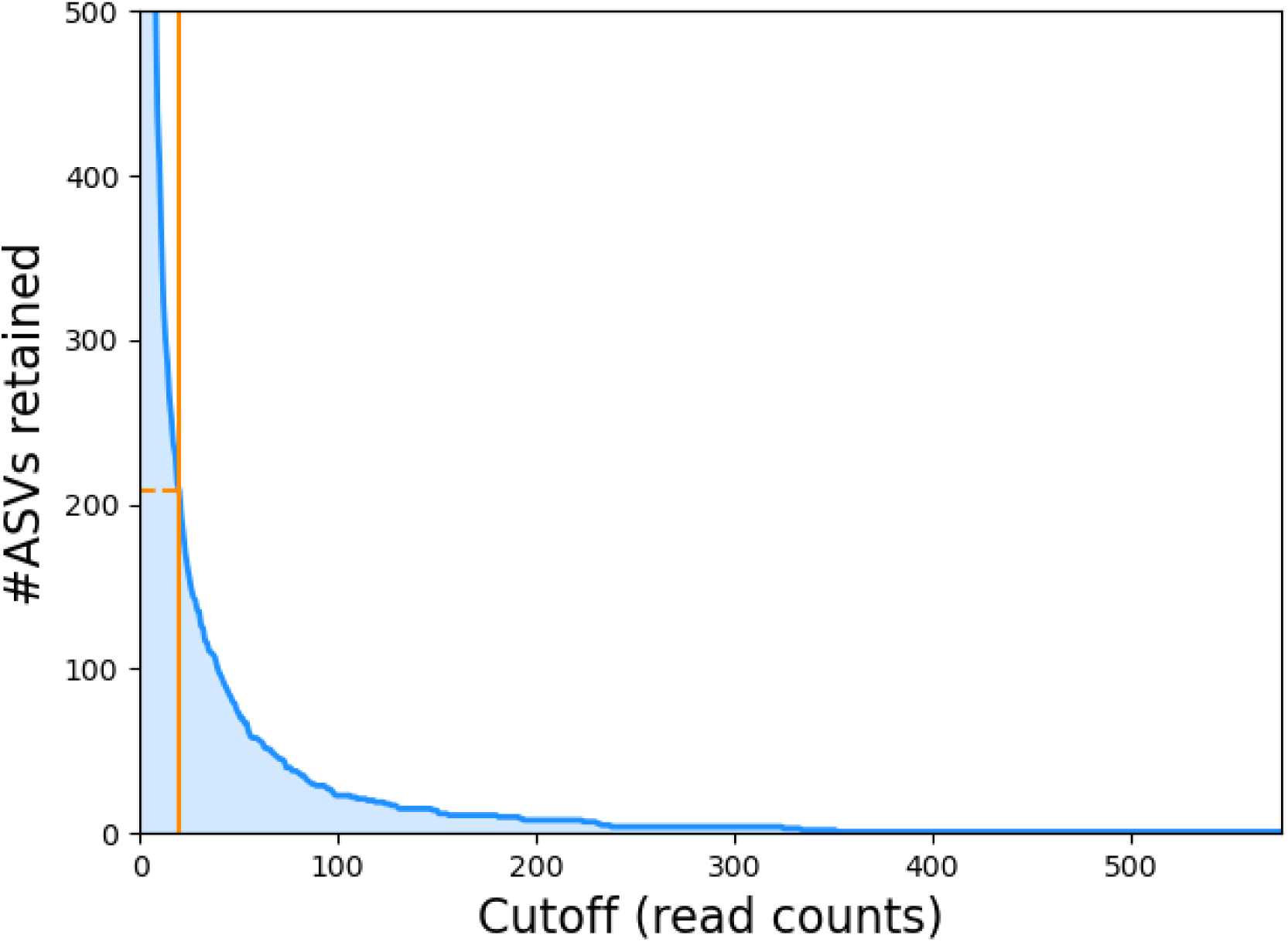
Number of ASVs retained with changing occurrence cutoff. Blue curve shows the number of ASVs retained with given cutoff used for selection. Orange line is the chosen cutoff (20 reads).

**Figure S7.**
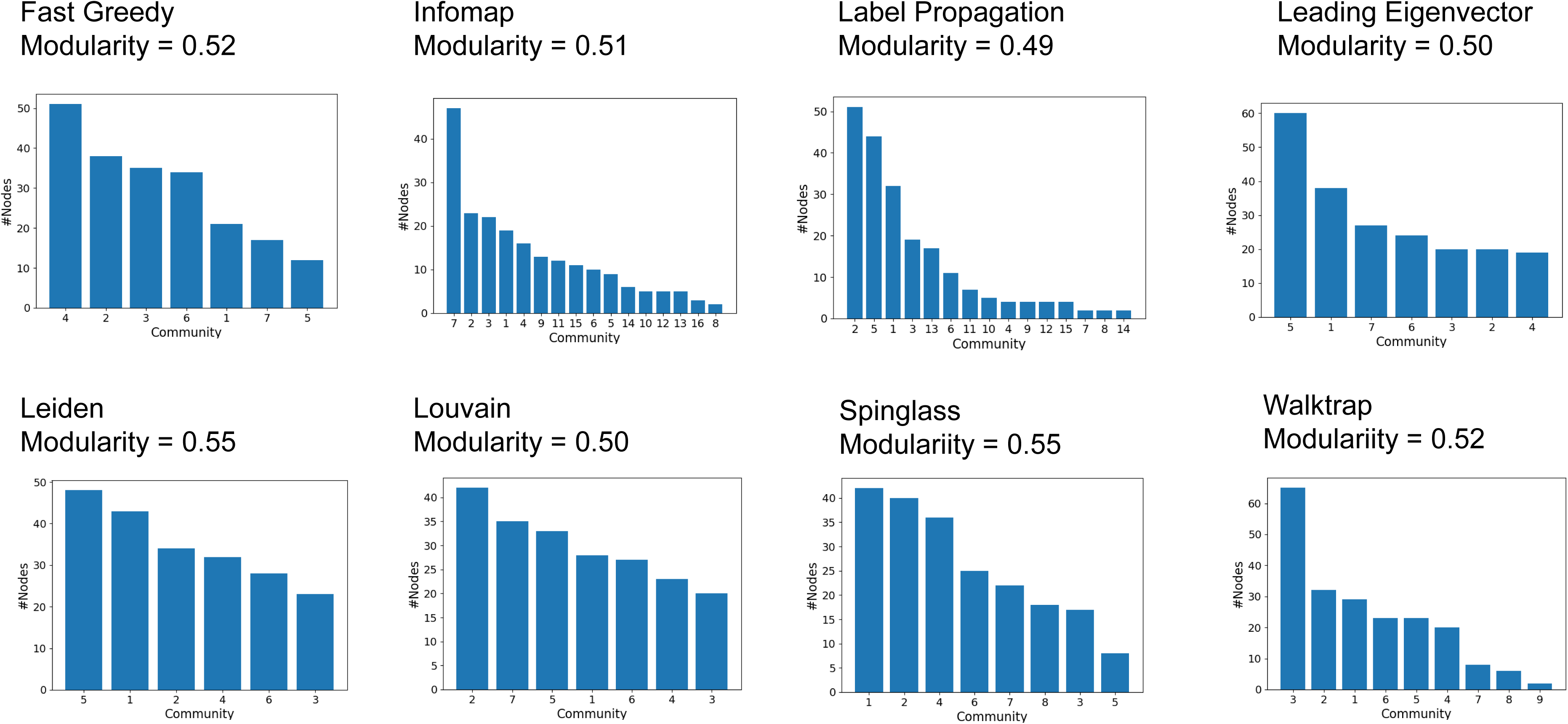
For each algorithm, modularity index and size of the communities. Modularity index of the community division by each algorithm and size of detected communities by it.

**Figure S8.**
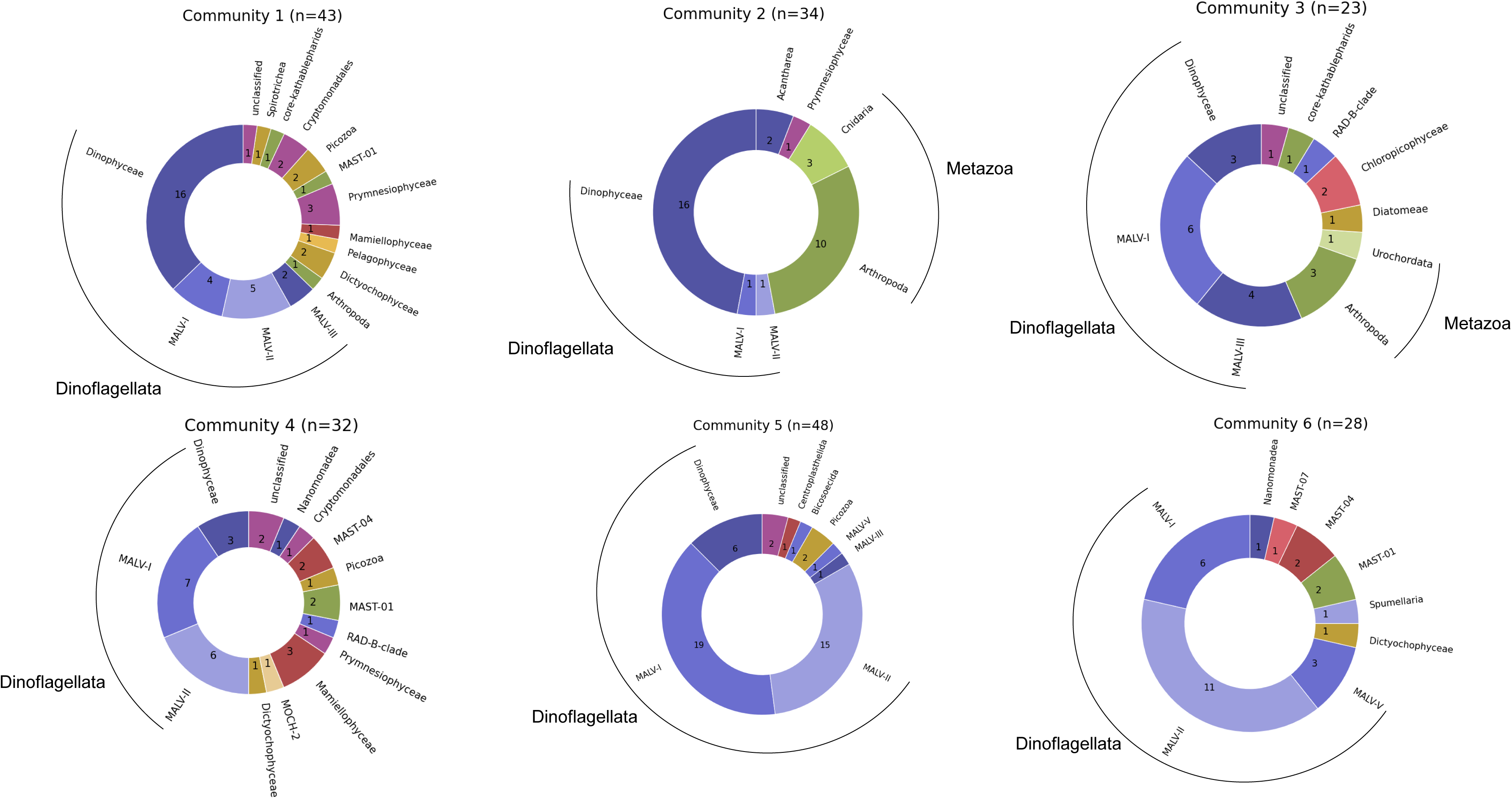
Taxonomic breakdown of plankton communities shown in circle charts. Taxonomic level is “taxogroup 2” in the EukRibo.

**Figure S9.**
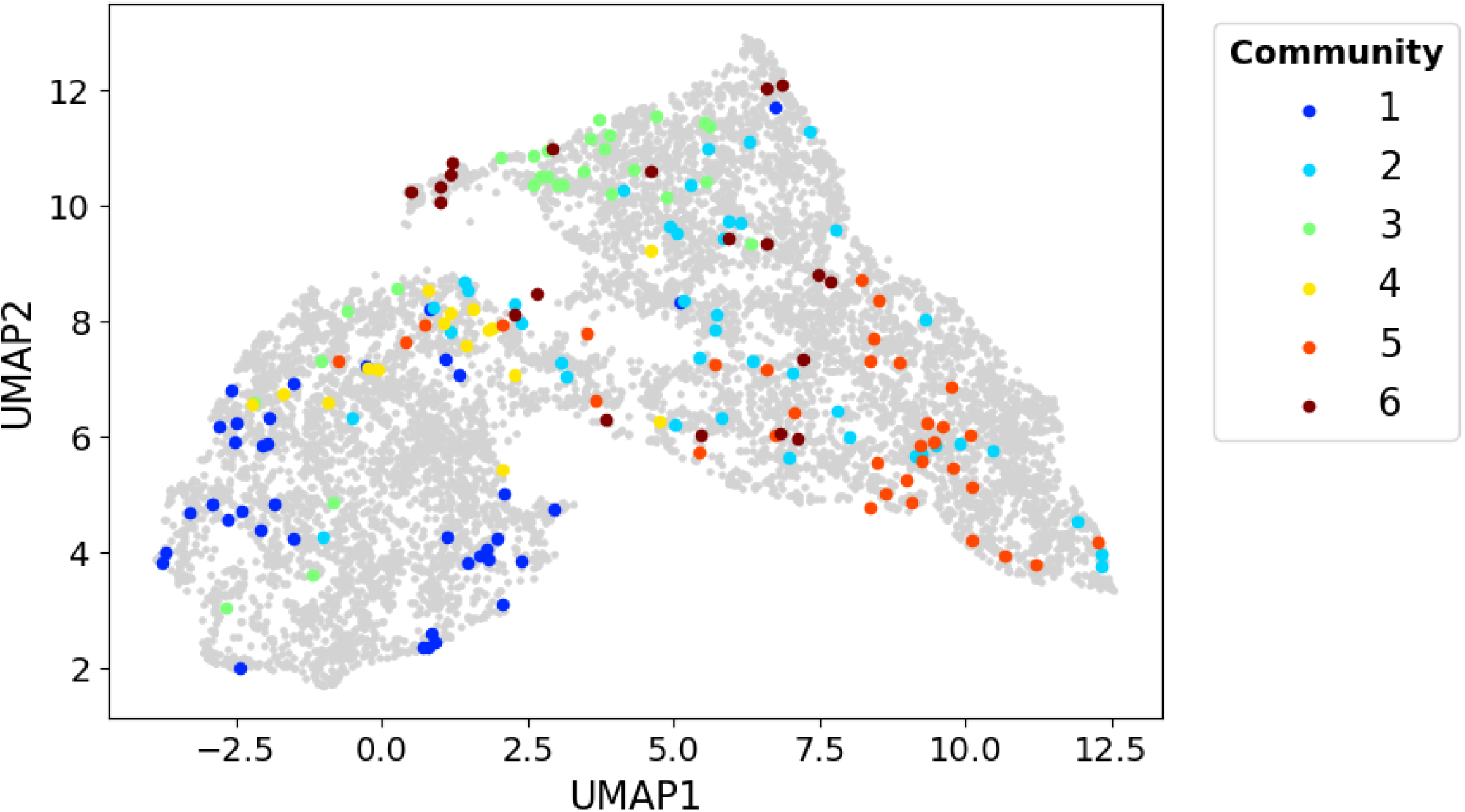
Representative community distribution in the satellite-derived parameter space. Metabarcoding samples projected on the 2-D map of the satellite-derived parameter space colored by representative community. Grey points are randomly selected grid cells used for learning the map.

**Figure S10.**
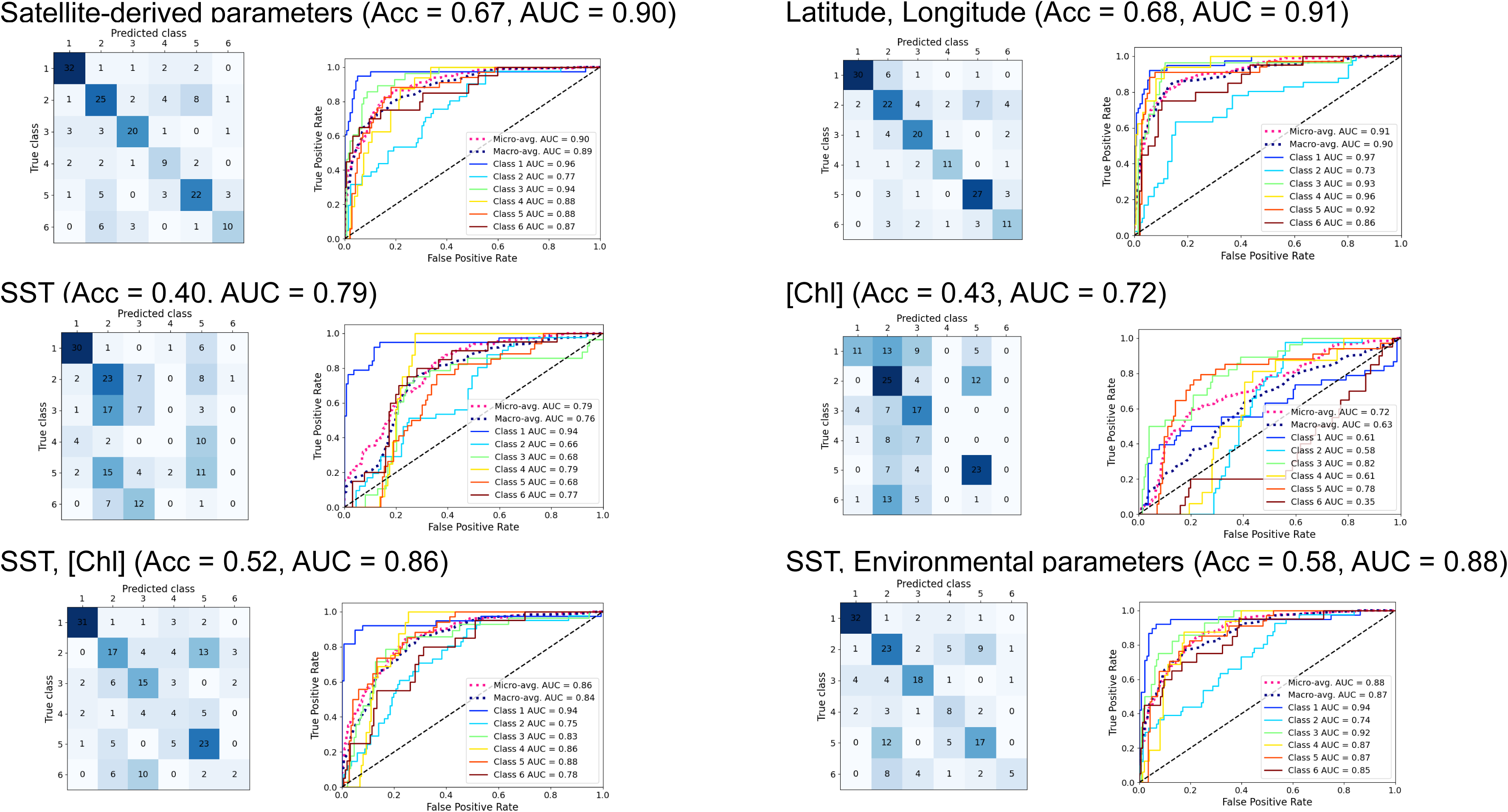
Comparison of prediction performance using different sets of satellite-derived and spatial parameters (leave-one-out cross-validation).

**Figure S11.**
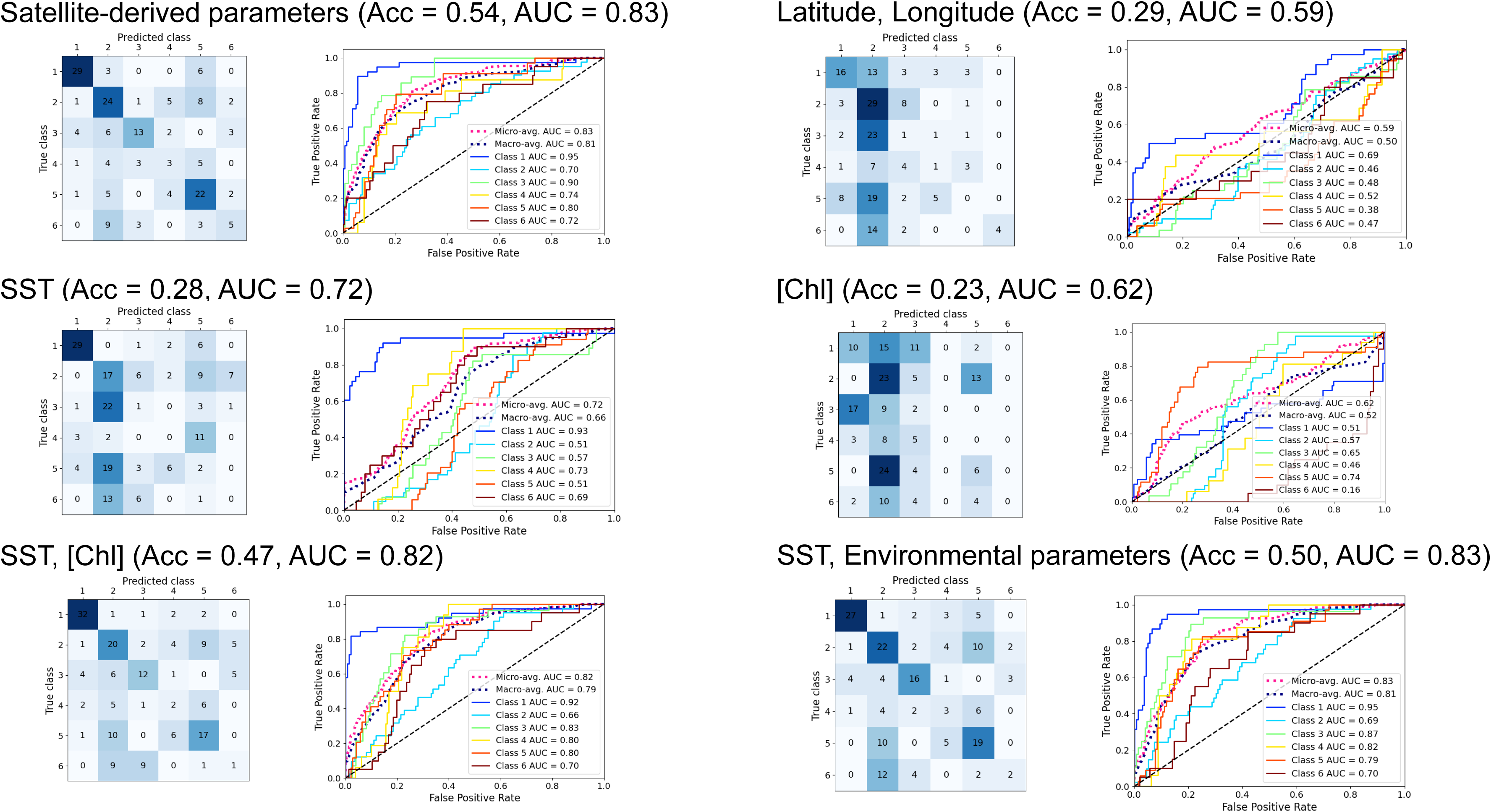
Comparison of prediction performance using different sets of satellite-derived and spatial parameters (buffered cross-validation).

**Figure S12.**
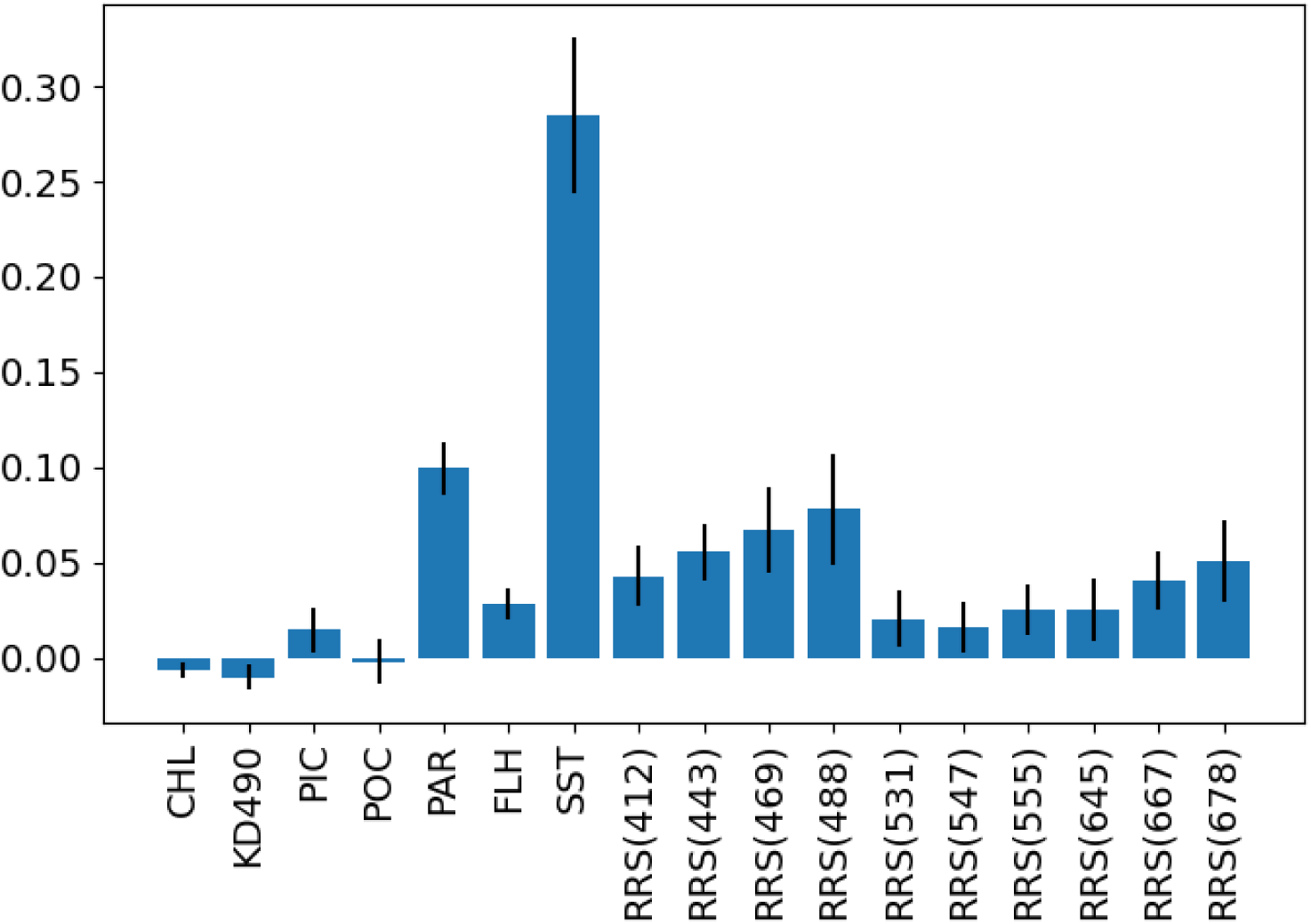
Permutation importance of each parameter in the full SVM model. Blue bars show mean of parameter importance over 5 times repeats. Error bars show standard deviation over repeats. CHL: chlorophyll *a* concentration, KD490: diffuse attenuation coefficient for downwelling irradiance at 490 nm, PIC/POC: particulate organic/inorganic carbon concentration, PAR: photosynthetically available radiation, FLH: normalized fluorescence line height, SST: sea surface temperature, RRS: remote sensing reflectance

**Figure S13.**
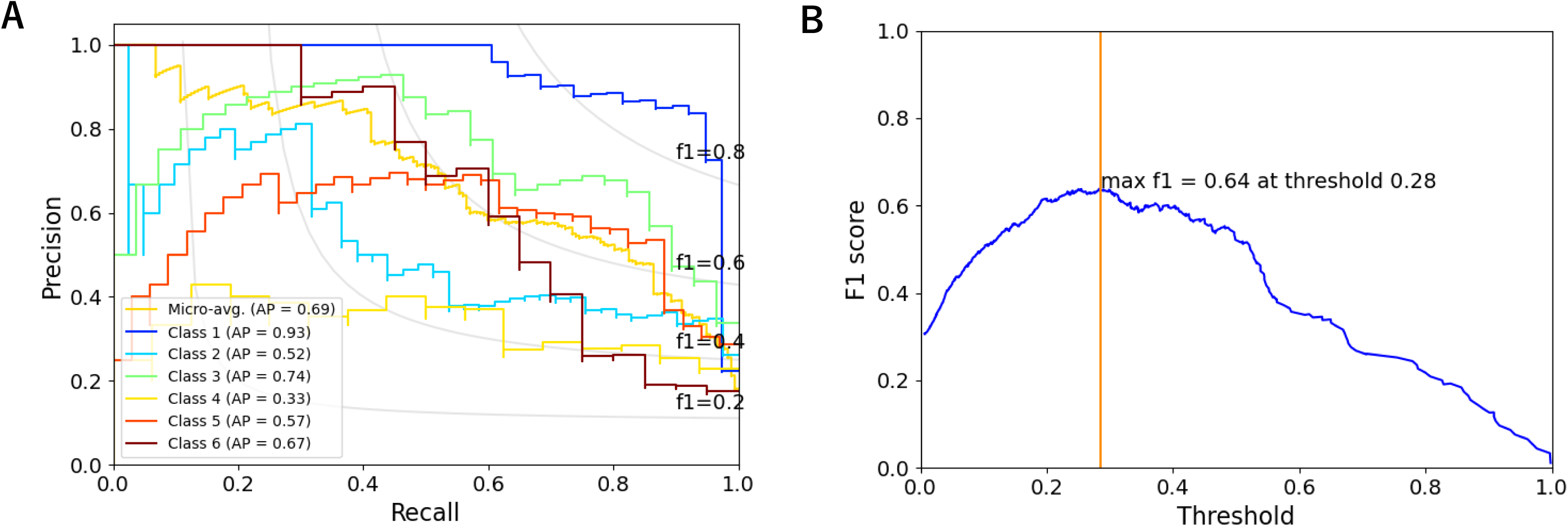
Precision, recall, and F1 score of SVM on community prediction based on satellite-devired parameters. **A** The precision-recall curve in the condition of leave-one-out cross-validation same as Figure 4B. **B** F1 score versus threshold of probabilistic output of SVM. Orange line shows the threshold making highest F1 score.

**Figure S14.**
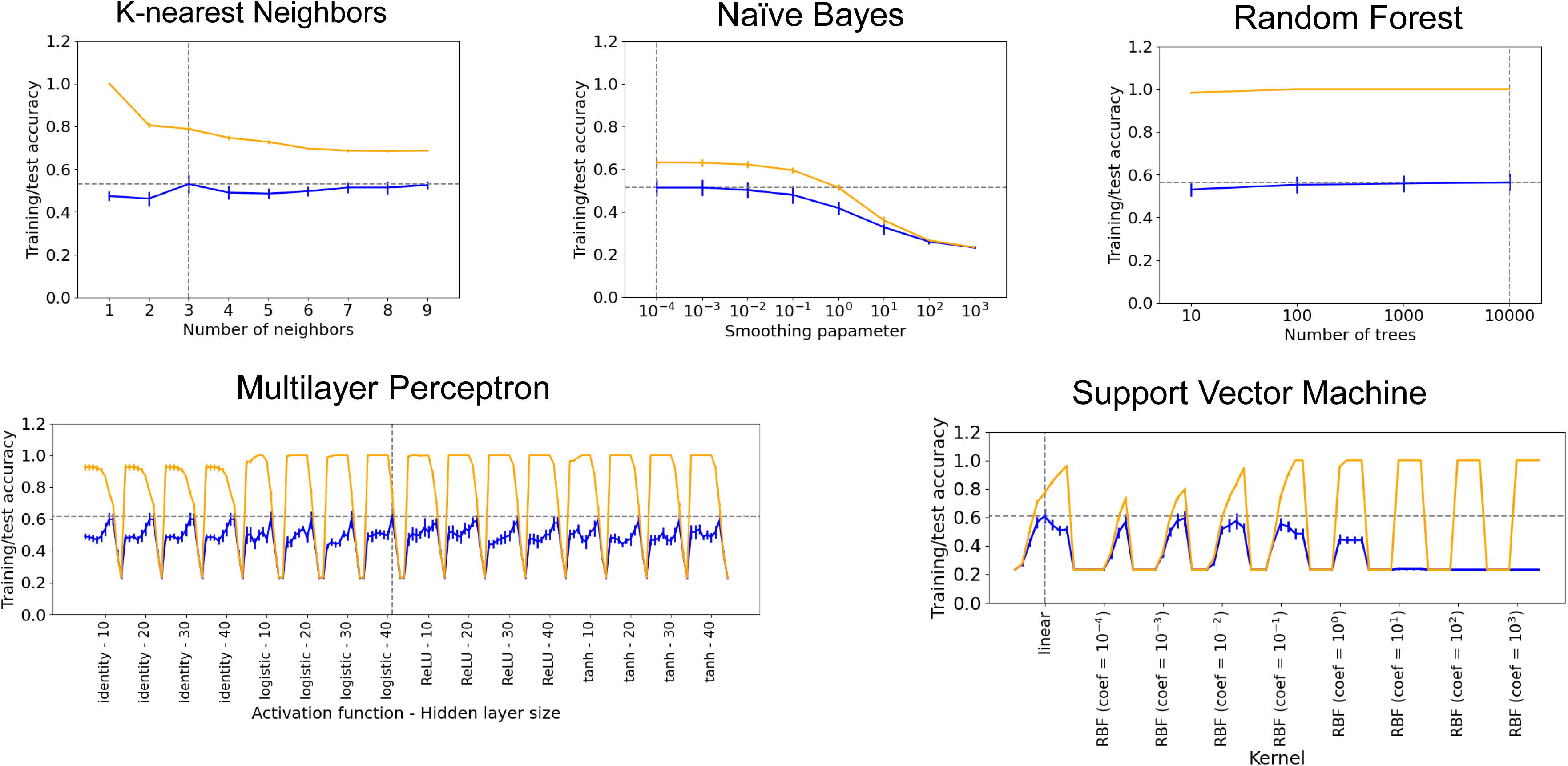
Grid search results in the training with all samples. Orange and blue lines show training and test accuracy, respectively. Gray dashed lines show the parameter with the best test accuracy. Ten L2 penalty parameters (10^-6^, 10^-5^, …, 10^3^; from left to right) were tested for each setting of Multilayer Perceptron and eight (10^-4^, 10^-3^, …, 10^3^) were tested for Support Vector Machine.

**Figure S15.**
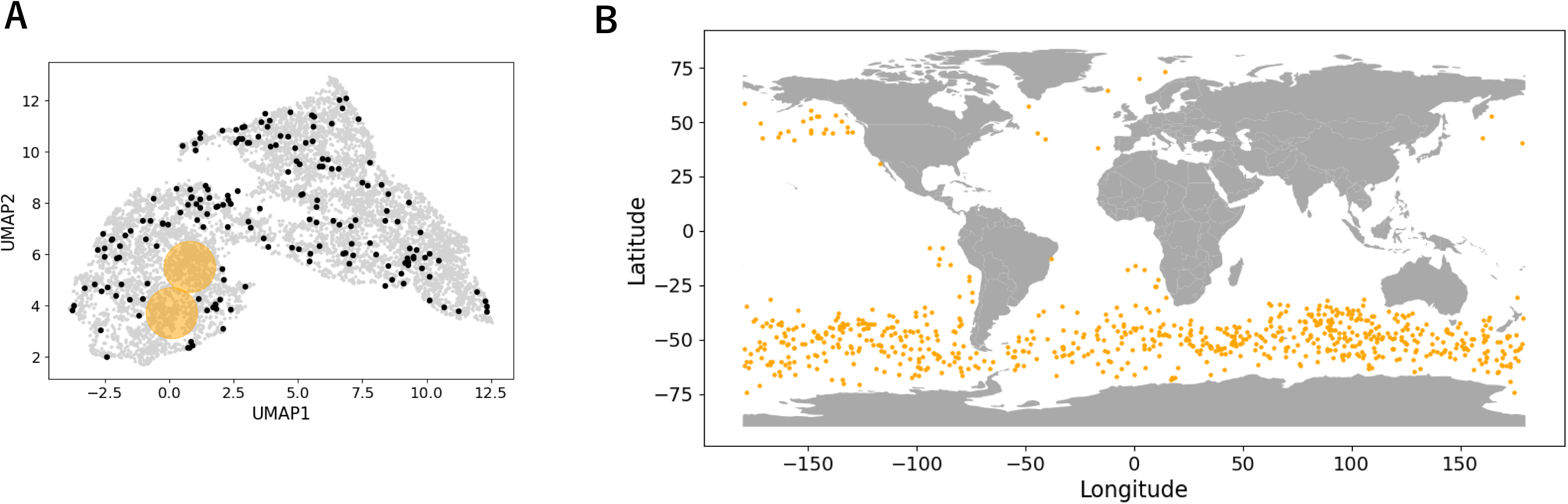
Location of samples in an unexplored region of the satellite-derived parameter space. **A** 2-D map of the satellite-derived parameter space. An unexplored region is shown in orange. **B** Geographic location of samples in the unexplored region of the parameter space (orange points).

**Figure S16.**
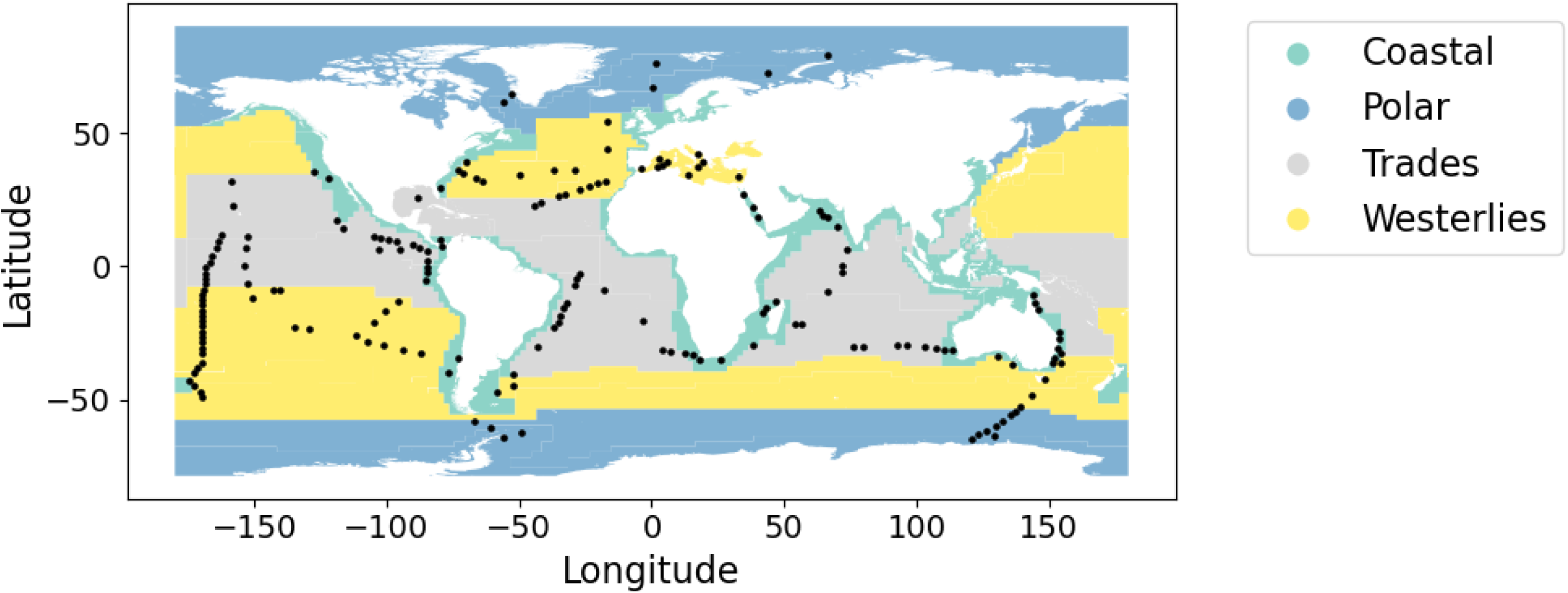
Map of Longhurst biome. Black points are metabarcoding samples. The shape file of the Longhurst biomes was downloaded from Marine Regions (https://www.marineregions.org).

